# Epidemiological inference for emerging viruses using segregating sites

**DOI:** 10.1101/2021.07.07.451508

**Authors:** Yeongseon Park, Michael Martin, Katia Koelle

**Affiliations:** Graduate Program in Population Biology, Ecology, and Evolution, Emory University, Atlanta, GA 30322; Department of Biology, Emory University, Atlanta, GA 30322; Emory Center of Excellence for Influenza Research and Response (CEIRR), Atlanta GA, USA

**Keywords:** statistical inference, segregating sites, viral sequence data, infectious disease modeling, phylodynamics, SARS-CoV-2

## Abstract

Epidemiological models are commonly fit to case data to estimate model parameters and to infer unobserved disease dynamics. Epidemiological models have also been fit to viral sequence data using phylodynamic inference approaches that rely on the reconstruction of viral phylogenies. However, especially early on in an expanding viral population, phylogenetic uncertainty can be substantial and methods that require integration over this uncertainty can be computationally intensive. Moreover, these approaches require estimation of parameters associated with models of sequence evolution in addition to the estimation of epidemiological parameters that are of primary interest. Here, we present an alternative approach to phylodynamic inference that can be used early on during the expansion of a viral lineage that circumvents the need for phylogenetic tree reconstruction. Our “tree-free” approach instead relies on quantifying the number of segregating sites observed in sets of sequences over time and using this trajectory of segregating sites to infer epidemiological parameters within a Sequential Monte Carlo (SMC) framework. Using forward simulations, we first show that epidemiological parameters and processes leave characteristic signatures in segregating site trajectories, demonstrating that these trajectories have the potential to be used for interfacing epidemiological models with sequence data. We then show that our proposed approach accurately recovers key epidemiological quantities such as the basic reproduction number and the timing of the index case from mock data simulated under a single-introduction scenario. Finally, we apply our approach to SARS-CoV-2 sequence data from France, estimating a reproductive number of approximately 2.5 to 2.7 under an epidemiological model structure that allows for multiple introductions, consistent with estimates from epidemiological surveillance data. Our findings indicate that “tree-free” statistical inference approaches that rely on simple population genetic summary statistics can be informative of epidemiological parameters and can be used for reconstructing infectious disease dynamics early on in an epidemic or during the expansion of a viral lineage.

## Introduction

Phylodynamic inference methods use viral sequence data to estimate epidemiological quantities such as the basic reproduction number and to reconstruct epidemiological patterns of incidence and prevalence. These inference methods have been applied to sequence data across a broad range of RNA viruses, including HIV (Stadler and Bonhoeffer 2013; Popinga et al. 2014; Ratmann et al. 2017; Volz et al. 2017), Ebola (Stadler et al. 2014; Vaughan et al. 2017; Volz and Siveroni 2018), dengue (Rasmussen et al. 2014), influenza (Rasmussen and Stadler), and most recently severe acute respiratory syndrome coronavirus 2 (SARS-CoV-2)(Danesh et al. 2020; Miller et al. 2020; Geidelberg et al. 2021). Most commonly, phylodynamic inference methods rely on underlying coalescent models or birth-death models. Coalescent-based approaches have been generalized to accommodate time-varying population sizes and structured epidemiological models, for example, susceptible-exposed-infected-recovered (SEIR) models and models with spatial subdivision (Volz 2012; Volz and Siveroni 2018). Birth-death approaches (Stadler 2010; Stadler et al. 2012), where a birth in the context of infectious diseases corresponds to a new infection and death corresponds to a recovery from infection, carry advantages such as capturing the role of demographic stochasticity in disease dynamics, which may be particularly important in emerging diseases that start with low infection numbers (Boskova et al. 2014). Both of these types of phylodynamic inference approaches rely on time- resolved phylogenies and have been incorporated into the phylogenetics software package BEAST2 (Bouckaert et al. 2014) to allow for joint estimation of epidemiological parameters and dynamics while integrating over phylogenetic uncertainty (Stadler et al. 2013; Volz and Siveroni 2018). Integrating over phylogenetic uncertainty is crucial when applying these methods to viral sequence data that are sampled over a short period of time and contain only low levels of genetic diversity. However, integrating over phylogenetic uncertainty can be computationally intensive. Moreover, phylodynamic approaches that use reconstructed trees for inference require estimation of parameters associated with models of sequence evolution, along with parameters that are of more immediate epidemiological interest.

Here, we present an alternative sequence-based statistical inference method that may be particularly useful when viral sequences are sampled over short time periods and when phylogenetic uncertainty present in time-resolved viral phylogenies is considerable. Instead of relying on viral phylogenies to infer epidemiological parameters or to reconstruct patterns of viral spread, the “tree-free” method we propose here fits epidemiological models to time series of the number of segregating sites (that is, the number of polymorphic sites) present in a sampled viral population. The approach we propose here allows for structured infectious disease models to be considered in a straightforward “plug-and-play” manner. It also incorporates the effect that demographic noise has on epidemiological dynamics. Below, we first describe how segregating site trajectories are calculated using sequence data and how they are impacted by sampling effort, rates of viral spread, and transmission heterogeneity. We then describe our proposed statistical inference method and apply it to simulated data to demonstrate the ability of this method to infer epidemiological parameters and to reconstruct unobserved epidemiological dynamics. Finally, we apply our segregating sites method to SARS-CoV-2 sequence data from France, arriving at quantitatively similar parameter estimates to those arrived at using epidemiological data.

### New Approaches

Mutations occur during viral replication within infected individuals and these have the potential to be transmitted. During the epidemiological spread of an emerging virus or viral lineage, the virus population (distributed across infected individuals) thus accrues mutations and diversifies genetically. This joint process of viral spread and evolution can be simulated forward in time using compartmental models, with patterns of epidemiological spread leaving signatures in the evolutionary trajectory of the virus population. Parameters of these compartmental models that govern patterns of epidemiological spread can thus in principle be estimated using viral sequence data. Here, we develop a statistical inference approach that fits compartmental epidemiological models to times series of a low-dimensional summary statistic calculated from sequence data. Specifically, we use trajectories of the number of segregating sites from samples of the viral population taken over time for statistical inference. In Materials and Methods, we provide details on the simulation of epidemiological models that incorporate viral evolution and thus can yield simulated time series of the number of segregating sites. In that section, we further describe our statistical inference approach that relies on using particle filtering (otherwise known as Sequential Monte Carlo; SMC) to infer parameters for these epidemiological models and to reconstruct unobserved disease dynamics. Because we propose the use of our method early on in an epidemic (or during the early expansion of a viral lineage), we focus primarily on estimating the basic reproduction number R_0_ using this inference approach.

## Results

### Segregating site trajectories are informative of epidemiological dynamics

The number of segregating sites present in a set of sampled viral sequences is defined as the number of nucleotide sites at which genetic variation is present in the sample set. To determine whether the number of segregating sites that are observed over time in a viral population may be informative of underlying epidemiological dynamics, we forward-simulated a classic susceptible-exposed-infected-recovered (SEIR) epidemiological model, augmented with viral evolution, under various sampling efforts and parameterizations (Figure 1; Materials and Methods). Simulations of this augmented SEIR model, initialized with a single infected individual, first indicate that segregating site trajectories are sensitive to sampling effort, as expected (Figure 1A,B). More specifically, we considered three different sampling strategies, each with sequences binned in non-overlapping 4-day time windows to calculate segregating site trajectories. These three sampling strategies consisted of a strategy with full sampling effort (all sequences per 4-day time window), one with dense sampling effort (40 sequences per 4-day time window) and one with sparse sampling effort (20 sequences per 4-day time window). With all three of these sampling efforts, the number of segregating sites first increases as the epidemic grows, with mutations accumulating in the virus population. Following the peak of the epidemic, the number of segregating sites starts to decline as viral sublineages die out, reducing the amount of genetic variation present in the viral population. A comparison between full, dense, and sparse sampling efforts indicates that lowering sampling effort results in a lower number of observed segregating sites during any time window. This is because at lower sampling effort, less of the genetic variation present in a viral population over a given time window is likely to be sampled. The patterns shown here across sampling strategies are robust to the time window length used for the calculation of segregating site trajectories (Figure S1).

**Figure 1.**
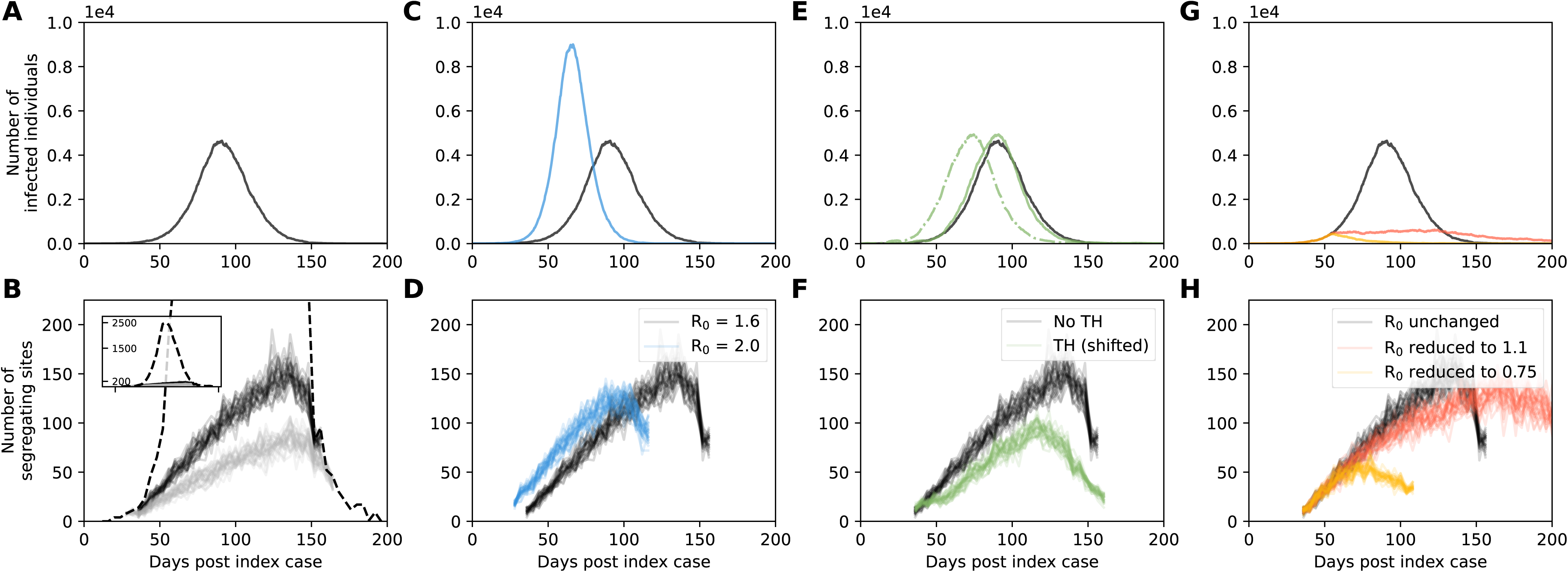
Segregating site trajectories under simulated epidemiological dynamics. (A) Simulated dynamics of infected individuals (I) under an SEIR model simulated with an R_0_ of 1.6. (B) Segregating site trajectories under full, dense, and sparse sampling efforts. The segregating site trajectory under full sampling effort is shown with a black dashed line. Dense sampling (black lines) corresponds to 40 sequences sampled per time window. Sparse sampling (gray lines) corresponds to 20 sequences sampled per time window. (C) Simulated dynamics of infected individuals (I) under an SEIR model simulated with an R_0_ of 2.0 (blue line) compared to those of the R_0_ = 1.6 simulation (black line). A higher transmission rate was used to generate the higher R_0_ value of 2.0. (D) Segregating site trajectories for the R_0_ = 2.0 simulation (blue lines) and the R_0_ = 1.6 simulation (black lines). (E) Simulated dynamics of infected individuals (I) under an SEIR model with an R_0_ of 1.6 and incorporating transmission heterogeneity (green, dashed line) compared to those of the original R_0_ = 1.6 simulation (black line) without transmission heterogeneity. Transmission heterogeneity was included by setting the parameter p_h_ to 0.06, resulting in 6% of the most infectious individuals being responsible for 80% of secondary infections. For ease of comparing segregating site trajectories, the transmission heterogeneity simulation was shifted later in time such that its epidemic peak aligned with the simulation without transmission heterogeneity (green, solid line). (F) Segregating site trajectories for the shifted transmission heterogeneity simulation (green lines) and the simulation without transmission heterogeneity (black lines). (G) Simulated dynamics of infected individuals (I) under an SEIR model simulated with changing R_0_. Changes in R_0_ occurred when the number of infected individuals reached 400. The simulation in red has R_0_ decreasing to 1.1. The simulation in yellow has R_0_ decreasing to 0.75. The simulation in black has R_0_ remaining at 1.6. (H) Segregating site trajectories for the three simulations shown in subplot G. Dense sampling effort (40 sequences sampled per time window) was used to generate all segregating site trajectories shown in subplots D, F, and G. 30 randomly-sampled segregating site trajectories are shown for each sampling effort in subplot B and for each epidemiological scenario in subplots D, F, and G. In all model simulations, y_E_ = 1/2 days , y_]_ = 1/3 days^-1^, population size N = 10^5^, and the per genome, per transmission mutation rate *μ* = 0.2. Initial conditions are S(t_0_) = N-1, E(t_0_) = 0, I(t_0_) = 1, and R(t_0_) = 0. For the transmission heterogeneity simulation (subplot E), initial conditions are S(t_0_) = N-1, E(t_0_) = 0, I_h_(t_0_) = 1, I_l_(t_0_) = 0, and R(t_0_) = 0. A time step of *τ* = 0.1 days was used in the Gillespie *τ* -leap algorithm.

To assess whether segregating site trajectories could be used for statistical inference, we first considered whether these trajectories differed between epidemics governed by different basic reproduction numbers (R_0_ values). Figure 1C shows simulations of the SEIR model under two parameterizations of the basic reproduction number: an R_0_ of 1.6, corresponding to the simulation shown in Figure 1A, and a higher R_0_ of 2.0. The epidemic with the higher R_0_ grew more rapidly (Figure 1C) and, under the same sampling effort, resulted in a more rapid increase in the number of segregating sites (Figure 1D). This indicates that segregating site trajectories can be informative of R_0_ early on in an epidemic.

We next considered the effect of transmission heterogeneity on segregating site trajectories. Many viral pathogens are characterized by ‘superspreading’ dynamics, where a relatively small proportion of infected individuals are responsible for a large proportion of secondary infections (Lloyd-Smith et al. 2005). The extent of transmission heterogeneity is often gauged relative to the 20/80 rule (the most infectious 20% of infected individuals are responsible for 80% of the secondary cases (Woolhouse et al. 1997)), with some pathogens like SARS-CoV-2 exhibiting extreme levels of superspreading, with as low as 10-15% of infected individuals responsible for 80% of secondary cases (Althouse et al. 2020; Miller et al. 2020; Lemieux et al. 2021; Sun et al. 2021). Because transmission heterogeneity is known to impact patterns of viral genetic diversity (Koelle and Rasmussen 2012), we simulated the above SEIR model with transmission heterogeneity to ascertain its effects on segregating site trajectories (Materials and Methods). Because transmission heterogeneity has a negligible impact on epidemiological dynamics once the number of infected individuals is large (Keeling and Rohani 2008), these simulated epidemiological dynamics should be quantitatively similar to one another, with transmission heterogeneity simply expected to shorten the timing of epidemic onset in simulations with successful invasion (Lloyd-Smith et al. 2005). Our simulations confirm this pattern (Figure 1E).

To compare segregating site trajectories between these simulations, we therefore shifted the simulation with transmission heterogeneity later in time such that the two simulated epidemics peaked at similar times (Figure 1E). Comparisons of segregating site trajectories between these simulations indicated that transmission heterogeneity decreases the number of segregating sites during any time window (Figure 1F). As expected, higher levels of transmission heterogeneity result in a greater decrease in the number of segregating sites (Figure S2). These results indicate that transmission heterogeneity needs to be taken into consideration when estimating epidemiological parameters using segregating site trajectories.

Finally, we wanted to assess whether changes in R_0_ over the course of an epidemic would leave signatures in segregating site trajectories. We considered this scenario because phylodynamic inference has often been used to quantify the effect of public health interventions on R_0_, most recently in the context of SARS-CoV-2 (Danesh et al. 2020; Miller et al. 2020). We thus implemented simulations with R_0_ starting at 1.6 and then either remaining at 1.6 or reduced to either 1.1 or 0.75 when the number of infected individuals reached 400 (Figure 1G). The segregating site trajectories for these three simulations indicate that reductions in R_0_ over the course of an epidemic leave signatures in this summary statistic of viral diversity (Figure 1H). The signatures left in the segregating site trajectories reflect the epidemiological dynamics that result from the reductions in R_0_. Reducing R_0_ to 1.1 results in a slower increase in the number of cases, a delayed epidemic peak, and a delayed decline in cases; as such, the number of segregating sites increases more slowly and peaks later. Reducing R_0_ to 0.75 results in an immediate decline in cases, with an observed drop in the number of segregating sites due to the stochastic loss of sublineages. Similar magnitude reductions in R_0_ that occurred later on in the simulated epidemic resulted in fainter signatures of this effect in the segregating site trajectories (Figure S3).

### Epidemiological inference using segregating site trajectories

To examine the extent to which inference based on segregating sites can be used for epidemiological parameter estimation, we generated a mock segregating site trajectory by forward simulating an SEIR model with a R_0_ of 1.6. From this simulation, we randomly sampled 500 viral sequences and binned these sequences into 4-day time windows based on their sampling times (Figure 2A). Figure 2B shows the segregating site trajectory from these binned sequences. From this trajectory, we first attempted to estimate only R_0_ under the assumption that the timing of the index case t_0_ is known. We estimated an R_0_ value of 1.54 (95% confidence interval of 1.37 to 1.81; Materials and Methods; Figure 2C), demonstrating that our segregating sites inference approach applied to this simulated dataset is able to recover the true R_0_ value of 1.6. Lower levels of sampling effort decreased the lower bound of the R_0_ estimate to 1.17 (Figure S4); the true R_0_ value of 1.6, however, still fell within the estimated 95% confidence interval. Instead of random sampling of infected individuals, adopting a more uniformly distributed sampling strategy acted to reduce the uncertainty in the R_0_ estimate (Figure S5). In Figure S6, we present results for the same set of sequences as those used in Figure 2, with the sequence data now binned in time windows of 1 day, 2 days, 6 days, and 10 days, instead of 4 days. These results show that R_0_ estimates are not biased by the use of different time window lengths.

**Figure 2.**
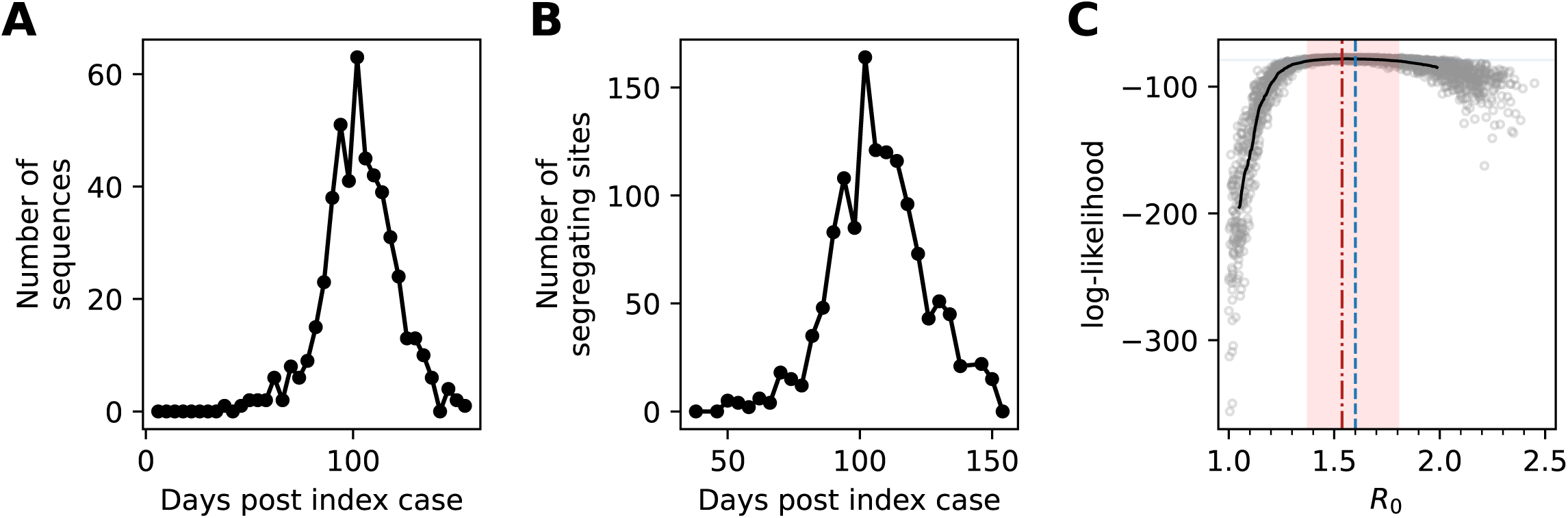
Epidemiological inference on a simulated trajectory of segregating sites. (A) The number of sampled sequences over time, binned by 4-day time windows. Sampling was done in proportion to the number of individuals recovering in a time window. In all, 500 sequences were sampled over the course of the simulated epidemic. (B) Simulated segregating site trajectory from the sampled sequences, by time window. (C) Estimation of R_0_ using Sequential Monte Carlo (SMC). Points show log-likelihood values from different SMC simulations with 2500 random drawn R_0_ values, uniformly sampled from a range between [1.0, 2.5). Solid black curve shows the moving average of 150 data points of the log-likelihood values. The vertical red dashed line shows the maximum likelihood estimate (MLE) of R_0_. The red band shows the 95% confidence interval of R_0_ (1.37-1.81). The vertical blue line shows the true value of R_0_ = 1.6. MLE and 95% CI were obtained from moving average log-likelihood values. Model parameters for the simulated data set are: R_O_= 1.6, y_E _= 1/2 days , y_]_ = 1/3 days , population size N = 10 , t_0_ = 0, and the per genome, per transmission mutation rate *μ* = 0.2. Initial conditions are S(t_0_) = N-1, E(t_0_) = 0, I(t_0_) = 1, and R(t_0_) = 0. A time step of *τ* = 0.1 days was used in the Gillespie *τ* -leap algorithm.

Because the timing of the index case (in cases with a single introduction) is almost certainly not known for an emerging epidemic, we further attempted to estimate both R_0_ and t_0_ using the segregating site trajectory shown in Figure 2B. To get an idea of the likelihood landscape over this parameter space, we first considered a range of R_0_ values between 1.0 to 2.5 and a range of t_0_ within 50 days of the true start date of 0. We then divided this parameter space into fine resolution parameter combinations (R_0_ intervals of 0.1 and t_0_ intervals of 2 days), and ran 20 SMC simulations for every parameter combination. In Figure 3A, we plot the mean value of the 20 SMC log-likelihoods for every parameter combination in the considered parameter space. Examination of this plot indicates that there is a log-likelihood ridge that runs between early t_0_/low R_0_ parameter combinations and late t_0_/high R_0_ parameter combinations. However, this ridge falls off on both sides, indicating that inference using segregating site trajectories can in principle estimate both t_0_ and R_0_. The maximum likelihood estimate for R_0_ was 1.7 (95% confidence = 1.5 to 2.0; true value = 1.6). The maximum likelihood estimate for t_0_ was 14 days (95% confidence = -12 to 32 days post the true t_0_ date of 0). Our results indicate that joint estimation of these parameters is thus possible in cases where a single introduction is responsible for igniting local circulation. Using our estimates of R_0_ and t_0_, we reconstructed the dynamics of the segregating sites (Figure 4A) and unobserved state variables: the number of susceptible, exposed, and infected individuals over time (Figures 4B, C, D). These reconstructed state variables captured the true epidemiological dynamics, demonstrating that our segregating sites approach can be used to infer epidemiological variables that generally go unobserved.

**Figure 3.**
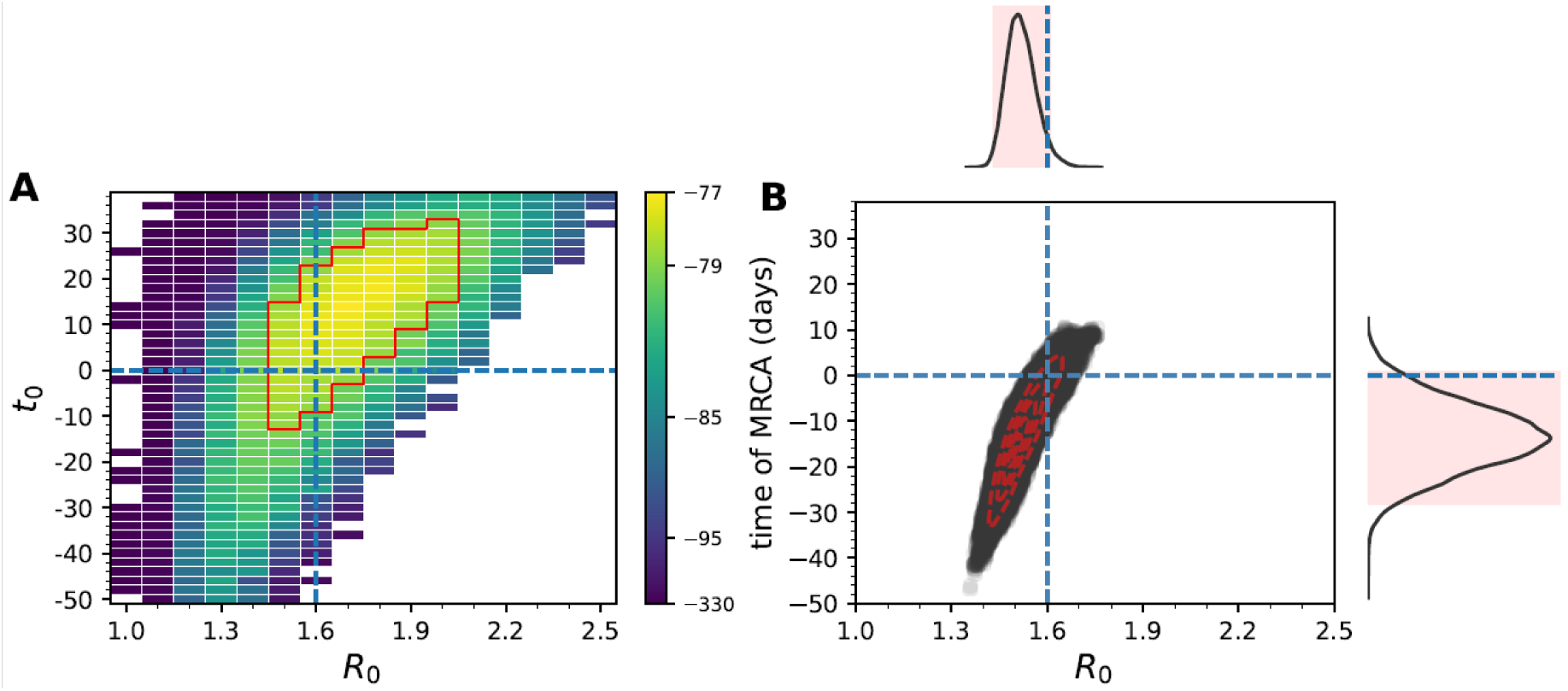
Joint estimation of the basic reproduction number (R_0_) and the timing of the index case (t_0_) using simulated data, and comparison against PhyDyn. (A) The likelihood surface based on the segregating site trajectory shown in Figure 2B is shown over a range of R_0_ and t_0_ values. Blank cells yielded log-likelihood values of negative infinity. The log-likelihood value shown in each cell is the mean log-likelihood value calculated from 20 SMC simulations. (B) Joint density plot for R_0_ and the time of the most recent common ancestor (tMRCA), as estimated using PhyDyn (Volz and Siveroni 2018) on the set of 500 sampled sequences shown in Figure 2A.

**Figure 4.**
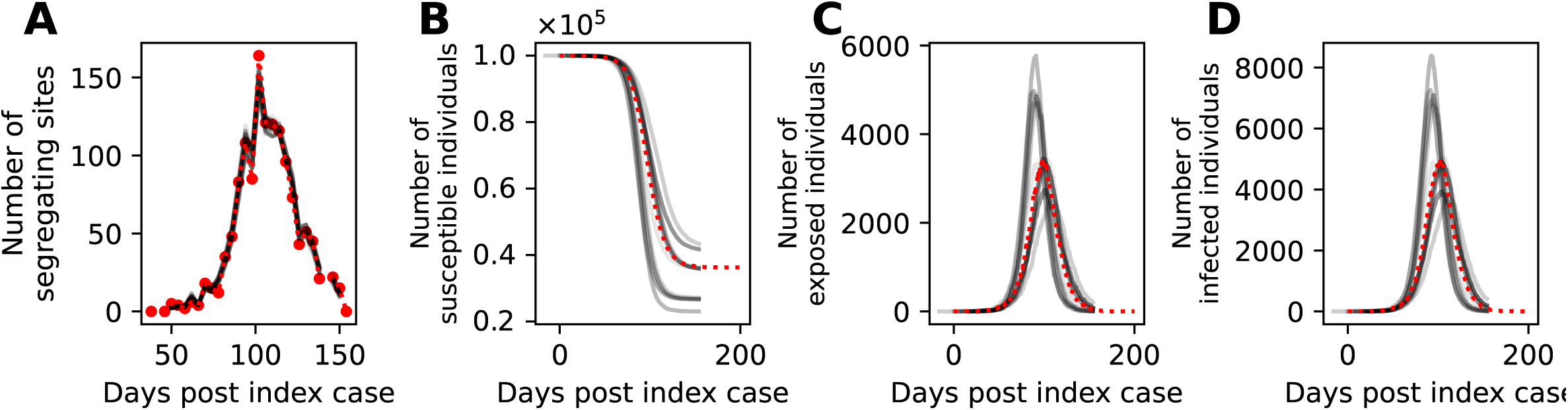
Reconstruction of unobserved state variables. (A) Simulated trajectory of the number of segregating sites (dashed red), alongside reconstructed trajectories of the number of segregating sites (gray). (B) Simulated dynamics of susceptible individuals (dashed red), alongside reconstructed dynamics of susceptible individuals (gray). (C) Simulated dynamics of exposed individuals (dashed red), alongside reconstructed dynamics of exposed individuals (gray). (D) Simulated dynamics of infected individuals (dashed red), alongside reconstructed dynamics of infected individuals (gray). Reconstructed state variables were obtained by running the particle filter using R_0_ and t_0_ parameter values sampled based on their mean log-likelihood values shown in Figure 3A.

As mentioned in the Introduction, there are existing phylodynamic inference approaches available that can estimate epidemiological model parameters using viral phylogenies that have been reconstructed from sequence data. Of particular note is the coalescent-based inference approach developed by Volz (Volz 2012) that has been implemented as PhyDyn (Volz and Siveroni 2018) in BEAST2. To compare our results using the segregating sites approach to results using PhyDyn, we generated mock viral nucleotide sequences from our set of 500 sampled sequences (Materials and Methods) and used these nucleotide sequences as input into PhyDyn. Assuming the same epidemiological model structure and using uninformative priors, PhyDyn was similarly able to recover the true R_0_ value of 1.6 used in the forward simulation (Figure 3B; 95% credible interval = 1.43 to 1.61). Because PhyDyn infers epidemiological parameters using a tree-based method, the program does not estimate the time of the index case t_0_. Instead, it estimates the time of the most recent common ancestor (tMRCA) of the viral phylogeny. This tMRCA is necessarily later than the time of the index case (Pekar et al. 2020). The tMRCA estimate’s credible interval spanned from -28.22 to 0.96 days post the true time of the index case (t_0_ = 0). As such, interpretation of the PhyDyn results would almost certainly result in timing the index case t_0_ as less than 0 (too early). The early estimate of t_0_ may be due to the “push-of-the-past” effect described by Nee and coauthors, which results from the assumption of deterministic dynamics in the inference process when the underlying population dynamics are stochastic (and conditioned on the persistence of a lineage)<u>(Nee et al. 1994)</u>. This “push-of-the-past” effect is usually reflected in an overestimate of the growth rate (or an overestimate in R_0_) in coalescent-based inference approaches that are applied to datasets with small population sizes (here, of infected individuals). Here, because R_0_ controls not only the rate of increase in the number of infected individuals at the start of the simulated pandemic but also the time at which the simulated pandemic starts to decline, the “push-of-the-past” effect may instead be reflected in a tMRCA estimate that occurs too early.

Because the impetus for developing the segregating sites inference approach was based on the extent of phylogenetic uncertainty present early on in an epidemic, we re-applied the inference approach to sequences sampled between day 32 and day 52 of the epidemic. Our results on this subset of simulated data indicate that R_0_ and t_0_ could again be jointly estimated, although the confidence intervals for R_0_ and t_0_ were both broader, as expected with a shorter time series (Figure 5A). Using this same set of sequences, PhyDyn estimated the credible interval for R_0_ to be 1.54 to 5.41, such that it tended to overestimate this parameter. In contrast to the PhyDyn results shown in Figure 3B, the tMRCA credible interval estimated by PhyDyn on this shorter time series was 14.5 to 31.0 days post the true t_0_ of 0 (Figure 5B), indicating potential consistency with this parameter. The tendency for PhyDyn to overestimate the basic reproduction number in this shorter time series that includes only the exponential growth phase of the simulated pandemic is consistent with expectations based on the “push-of-the-past” effect.

**Figure 5.**
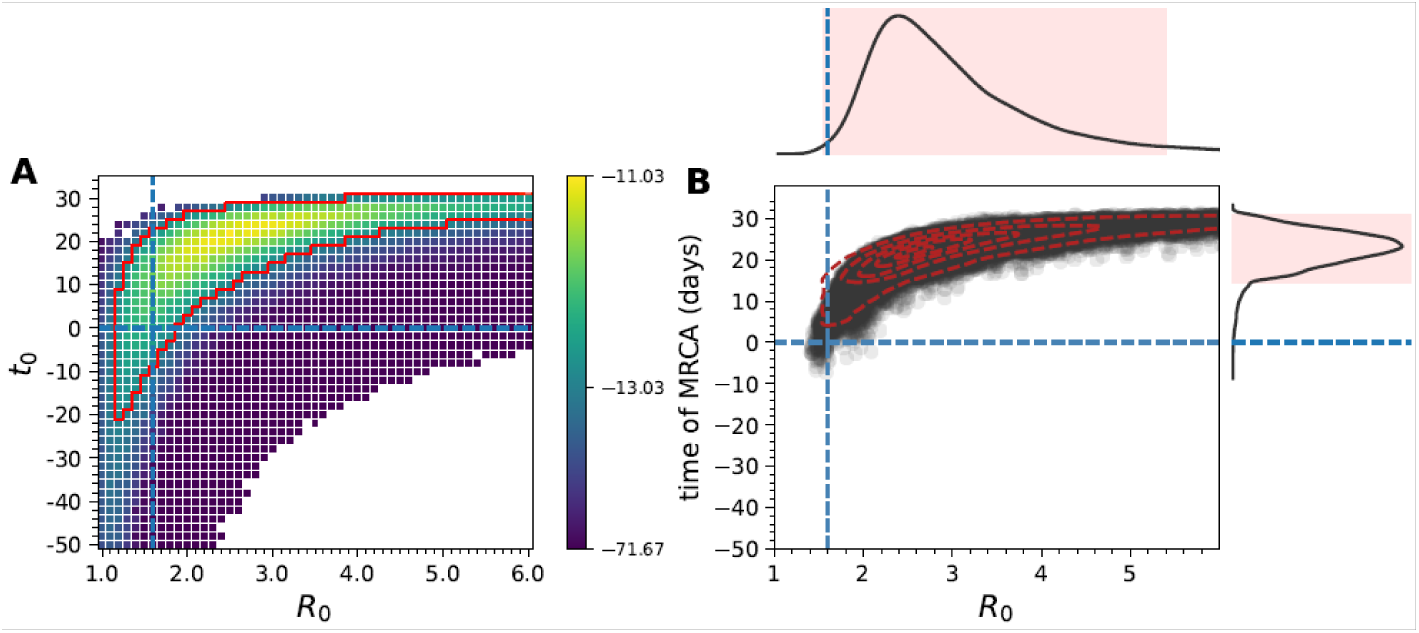
Joint estimation of the basic reproduction number (R_0_) and the timing of the index case (t_0_) using early samples from the simulation, and comparison against PhyDyn. (A) The likelihood surface based on a segregating site trajectory calculated using sequences from day 32 through day 52 of the simulated pandemic. Sequences were binned into 4-day windows, with 10 individuals sampled from each window. Blank cells yielded log-likelihood values of negative infinity. As in Figure 3A, the log- likelihood value shown in each cell is the mean log-likelihood value calculated from 20 SMC simulations. (B) Joint density plot for R_0_ and the time of the most recent common ancestor (tMRCA), as estimated using PhyDyn on the same set of 50 sampled sequences.

### Epidemiological inference using SARS-CoV-2 sequences from France

We applied the segregating sites inference approach to a set of SARS-CoV-2 sequences sampled from France between January 23, 2020 and March 17, 2020, when a country-wide lockdown was implemented. We decided to apply our approach to this set of sequences for several reasons. First, a large fraction of the 479 available full-genome sequences from France over this time period appear to be genetically very similar to one another (Gámbaro et al. 2020), indicating that one major lineage may have taken off in France (or at least, that most samples stemmed from one major lineage). This lineage would be the focus of our analysis. Second, an in-depth epidemiological analysis previously inferred R_0_ for France prior to the March 17 lock- down measures that were implemented (Salje et al. 2020). This analysis fit a compartmental infectious disease model to epidemiological data that included case, hospitalization, and death data. Because our segregating sites inference approach can accommodate epidemiological model structures of arbitrary complexity, we can adopt the same model structure as in this previous analysis. We can also set the epidemiological parameters that are assumed fixed in this previous analysis to their same values. By controlling for model structure and the set of model parameters assumed as given, we can ask to what extent sequence data corroborate the R_0_ estimates arrived at from detailed fits to epidemiological data.

To apply our segregating sites approach to the viral sequences from France, we first identified the subset of the 479 sequences that constituted a single, large lineage. To keep with the “tree-free” emphasis of our approach, we identified this subset of n = 432 sequences without inferring a phylogeny (Materials and Methods). Using phylogenetic inference, however, we confirmed that our subset of sequences constituted a single clade, with excluded sequences from France falling outside of this clade (Figure S7). To generate a segregating site trajectory from these sequences, we defined 4-day time windows such that the last time window ended on March 17, 2020. Figure 6A shows the number of sequences falling into each time window. Figure 6B shows the segregating site trajectory calculated from these sequences.

**Figure 6.**
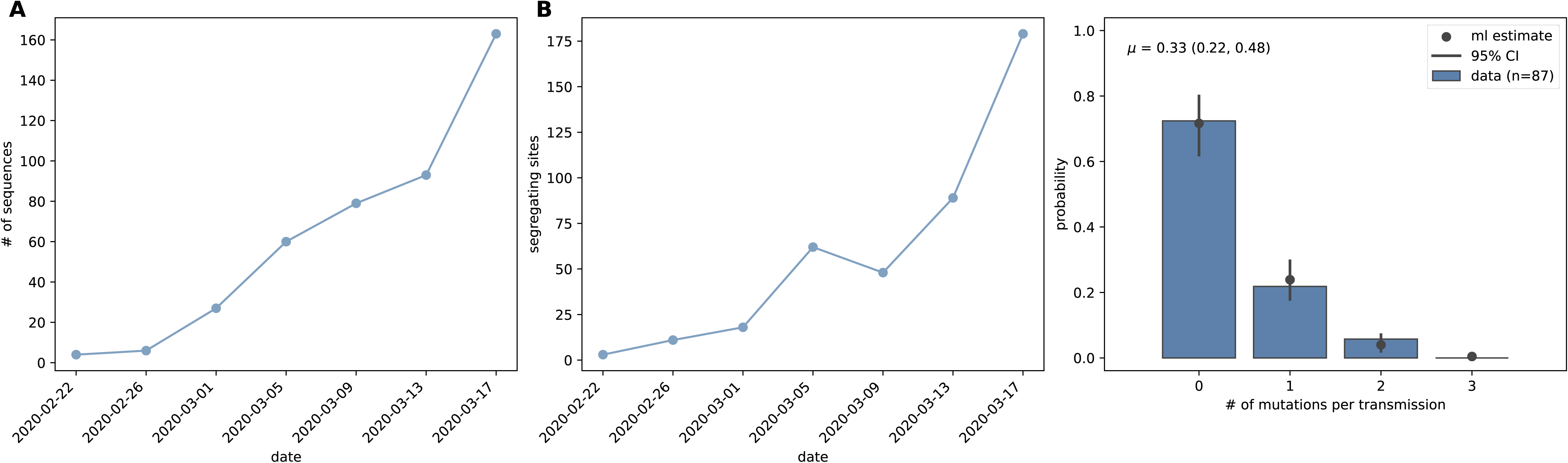
Sequences and parameters used for epidemiological inference based on SARS-CoV-2 sequences from France. (A) The number of sampled sequences over time, calculated using a 4-day time window. (B) The segregating site trajectory calculated from the binned sequences shown in (A). (C) Estimation of the per-genome, per-transmission mutation rate *μ*. The histogram shows the fraction of transmission pairs with consensus sequences that differed from one another by the number of mutations shown on the x-axis. The Poisson estimate of the mutation rate from these data (black) is *μ* = 0.33 (95% CI = 0.22-0.48).

We parameterized the model with a per genome, per transmission mutation rate of *μ* = 0.33 using consensus sequence data from established SARS-CoV-2 transmission pairs that were available in the literature (James et al. 2020; Popa et al. 2020; Braun et al. 2021; Lythgoe et al. 2021) (Materials and Methods). Specifically, for each of the 87 transmission pairs we had access to, we calculated the nucleotide distance between the consensus sequence of the donor sample and that of the recipient sample and fit a Poisson distribution to these data (Figure 6C). Using this approach, we estimated a *μ* value of 0.33 (95% confidence interval of 0.22 to 0.48), corresponding approximately to one mutation occurring every 3 transmission events.

Similar to the approach we undertook with our simulated data, we first attempted to estimate R_0_ and the timing of the index case t_0_ for this segregating site trajectory. We considered a broad parameter space over which to calculate log-likelihood values. Specifically, we considered R_0_ values between 1.0 and 4.5 and t_0_ values of December 1^st^, 2019 and February 14^th^, 2020. We ran 20 SMC simulations and calculated the mean log-likelihood for each parameter combination (Figure 7). We estimated R_0_ to be 2.3 (95% confidence interval = 1.7 to 3.6), consistent with the R_0_ estimate of 2.9 (95% confidence interval = 2.81 to 3.01) arrived at through epidemiological time series analysis (Salje et al. 2020). The maximum likelihood estimated of t_0_ was January 18^th^, 2020 (95% confidence interval = December 22^nd^ 2019 to February 8^th^, 2020). Our maximum likelihood estimate of t_0_ in the middle of January 2020 aligns well with findings from Gámbaro et al. (2020).

**Figure 7.**
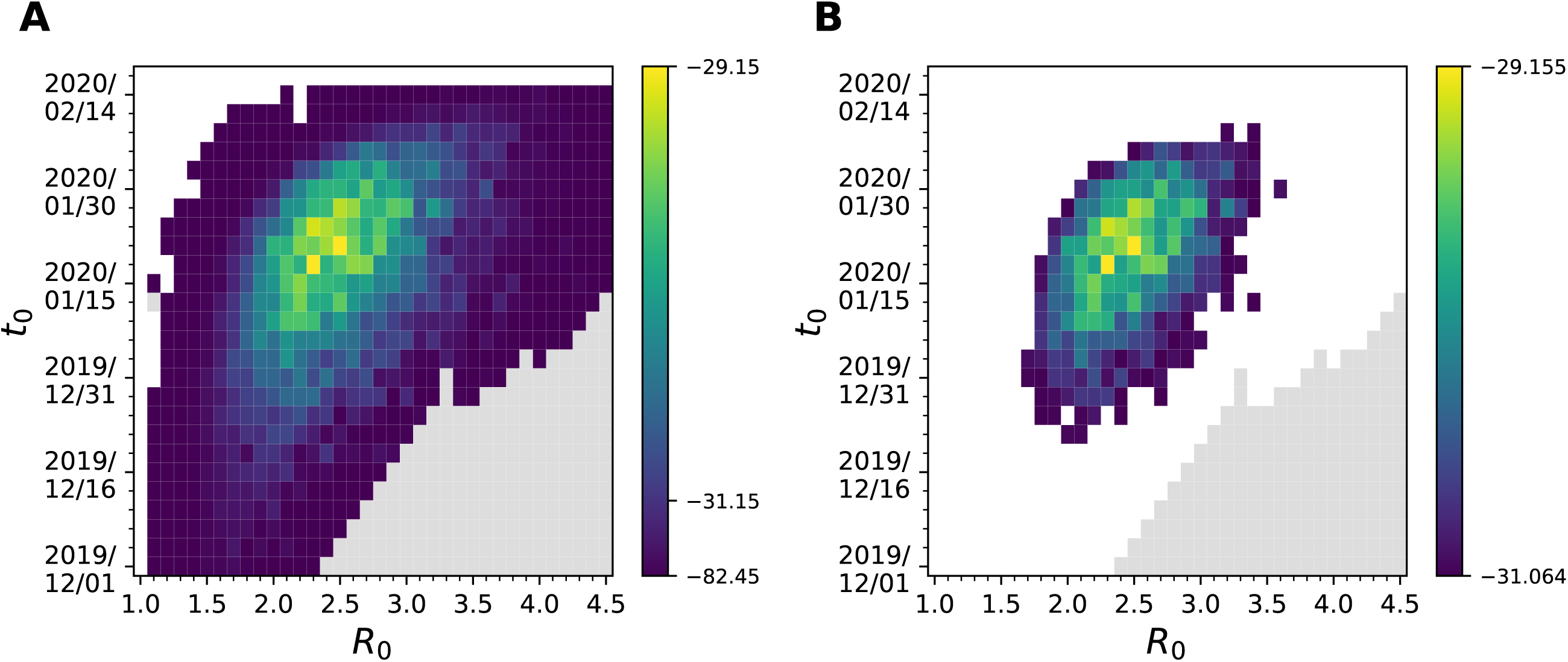
Joint estimation of the basic reproduction number R_0_ and the time of the index case t_0_ for the France SARS-CoV-2 data. (A) The joint log-likelihood surface based on the estimated segregating site trajectory for the France data. Each cell shows the mean log-likelihood value based on 20 SMC simulations. Blank cells indicate mean log-likelihood values of negative infinity. Gray cells indicate where log-likelihood values were not evaluated due to extended simulation time. (B) The same joint log- likelihood surface showing cells as colored only if they fell within the 95% confidence interval of the parameter estimates.

Despite the reasonable estimate for R_0_ we obtained under this model, we decided to also consider an alternative model that allowed for multiple introductions of the focal lineage into France (Materials and Methods). This decision was based on evolutionary analyses that have shown that regional SARS-CoV-2 epidemics in Europe (as well as in the US) were initiated through multiple introductions rather than only a single one (Worobey et al. 2020). Instead of attempting to estimate R_0_ and t_0_, we attempted to jointly estimate R_0_ and a parameter *η* using the segregating site trajectory. The parameter *η* quantifies the extent to which transmission between France and regions outside of France is reduced relative to transmission occurring within France. This model further required specification of the time at which the basal genotype evolved in the outside reservoir, which we refer to as t_e_. We considered a broad parameter space over which to calculate log-likelihood values (R_0_ values between 1.0 and 4.0 and *η* values between 10^-8^ and 10^-2^) and three different t_e_ values: December 24^th^, 2019, January 1^st^, 2020, and January 8^th^, 2020 (Materials and Methods). At each of these t_e_ values, we ran 20 SMC simulations and calculated the mean log-likelihood for each parameter combination (Figure 8A-C). The maximum likelihood estimates of R_0_ were 2.7 (95% confidence interval = 2.1 to 3.6), 2.5 (95% confidence interval = 2.0 to 3.7), and 2.7 (95% confidence interval = 1.9 to 3.7), respectively, under t_e_ = December 24^th^, 2019, January 1^st^, 2020 and January 8^th^, 2020. These results indicate that the inferred R_0_ values are relatively insensitive to the assumed emergence time of the basal genotype in the outside reservoir. At later emergence times, the estimates for *η* were higher, indicating that a later emergence time was compensated for by a higher transmission rate between the outside reservoir and individuals in France.

**Figure 8.**
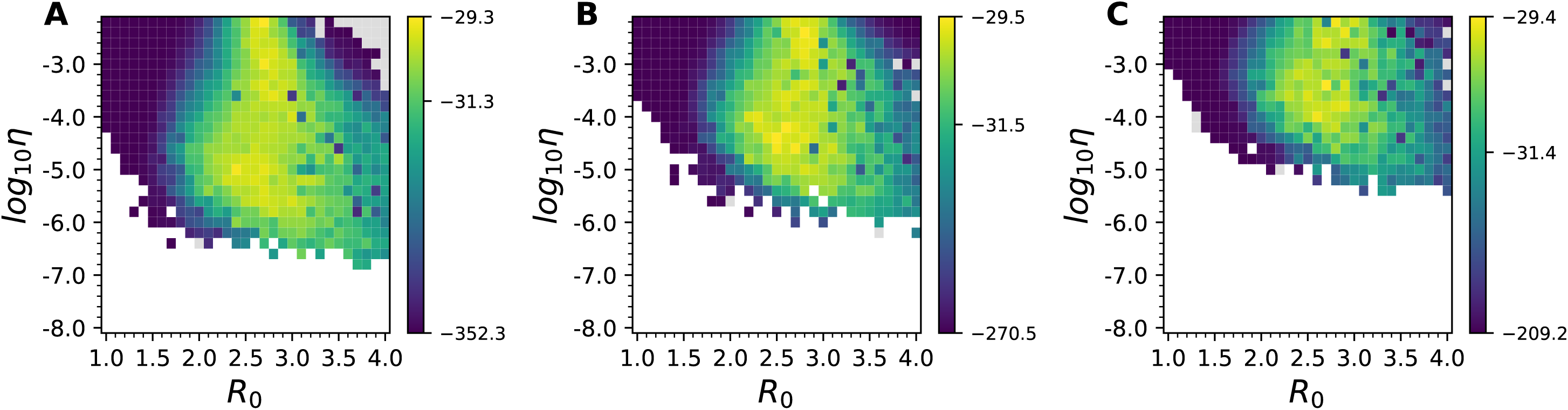
Joint estimation of the basic reproduction number R_0_ and the transmission-reduction parameter *η* for the multiple-introductions model using the France data. The joint log-likelihood surface based on the estimated segregating site trajectory for the France data is shown under three different basal genotype emergence times: t_e_ = December 24, 2019 (A), January 1, 2020 (B), and January 8, 2020 (C). Each cell shows the mean log-likelihood value based on 20 SMC simulations. Blank cells indicate mean log-likelihood values of negative infinity. Gray cells indicate where log-likelihood values were not evaluated due to extended simulation time.

We reconstructed the unobserved state variables for the multiple-introductions model using SMC simulations parameterized with R_0_ and *η* values that were sampled from the parameter space shown in Figure 8, according to the same approach as the one we used for reconstructing state variables on the mock segregating sites trajectory (Figure 9). As expected for an epidemic with an R_0_ > 1, the total number of infected individuals increased exponentially over the time period considered (Figures 9D-F). In Figures 9G-I, we plot the reconstructed cumulative number of recovered individuals over time. These cumulative trajectories indicate that by mid-March 2020, approximately 0.02% to 0.15% of individuals in France had recovered from infection from this SARS-CoV-2 lineage. These cumulative predictions can be compared against findings from a serological study that was conducted over this time period in France (Le Vu et al. 2021). Based on a survey of 3221 individuals, this study found that 0.41% of individuals (95% confidence interval = 0.05% to 0.88%) had gotten infected with SARS-CoV-2 by March 9 to 15, 2020 (Figures 9G-I). Our estimates fall in line with these independent estimates. Finally, in Figures 9J-L, we plotted the reconstructed cumulative number of infections that resulted directly from contact with individuals outside of France. By the first sampled time window (ending on February 22, 2022), the sampled particles showed evidence of inferred repeated introductions of this lineage into France, with the majority of particles pointing towards 5-100 introductions of these lineage into France prior to February 22.

**Figure 9.**
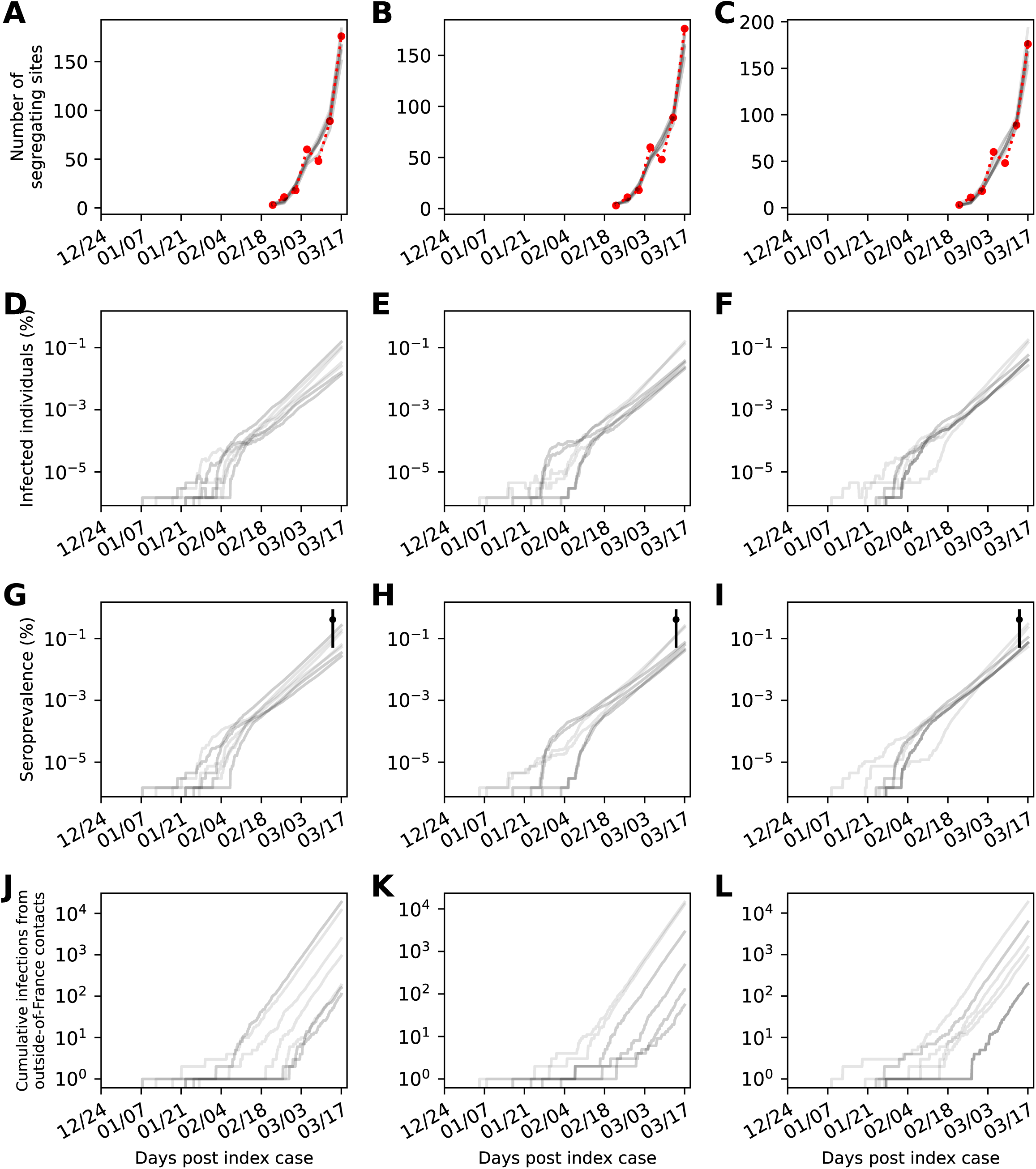
Trajectories of reconstructed state variables for the France data under the multiple- introductions model. State variables are reconstructed for the multiple-introductions model with three different values assumed for the emergence time of the basal genotype: t_e_ = December 24, 2019 (first column), January 1, 2020 (second column), and January 8, 2020 (third column). (A-C) Segregating site trajectory for the France SARS-CoV-2 data (ref), alongside segregating site trajectories from 10 sampled SMC particles (gray). (D-F) Reconstructed dynamics of the total number of infected individuals (E_1_ + E_2_ + I), shown in percent of France’s population. (G-I) Reconstructed dynamics of the cumulative number of recovered individuals over time, shown in percent of France’s population. Independent estimates of the fraction of the population that has been infected with SARS-CoV-2 by mid-March are shown in black. Estimates are from a serological study conducted during the time window March 9-15, 2020 (Le Vu et al. 2021). (J-L) Reconstructed dynamics of the cumulative number of infections in France that resulted from contact with infected individuals outside of France. Reconstructed state variables shown in panels (A-L) are based on 10 SMC simulations using parameter combinations of R_0_ and *η* that were sampled in proportion to their likelihood values.

## Discussion

Here, we developed a statistical inference approach to estimate epidemiological parameters from virus sequence data. Our inference approach is a “tree-free” approach in that it does not rely on the reconstruction of viral phylogenies to estimate model parameters. One benefit of using such an approach for parameter estimation of emerging viral pathogens is that, early on in an epidemic, phylogenetic uncertainty present in time-resolved viral phylogenies is significant, and tree-based phylodynamic inference approaches would need to integrate over this uncertainty. This is often times computationally intensive. A second benefit of using a tree-free approach is that parameters of the model of sequence evolution do not need to be estimated, reducing degrees of freedom considerably. Instead of viral phylogenies being the data that statistically interface with the epidemiological models, our approach uses a population genetic summary statistic of the sequence data, namely the number of segregating sites present in time-binned sets of viral sequences. Our inference approach benefits from being plug-and-play in that it can easily accommodate different epidemiological model structures. Based on fits to a simulated data set, we have shown that segregating site trajectories can be used to estimate important epidemiological parameters such as R_0_ and the timing of the index case t_0_ in cases where a single introduction can be assumed. We further fit a multiple-introductions epidemiological model to a segregating site trajectory that was calculated from SARS-CoV-2 sequence data from France, estimating a basic reproduction number R_0_ of approximately 2.5-2.7 (with a 95% confidence interval of approximately 2.0 to 3.7). These results are consistent with previous estimates from an epidemiological analysis and consistent with a serological study conducted in mid-March 2020.

Our inference approach relies on several assumptions. Most notably, it relies on an assumption of random sampling of individuals and on an assumption that all genetic variation that is observed is phenotypically neutral. These assumptions are shared by existing phylodynamic inference methods, although some newer statistical developments have relaxed the assumption of neutrality by incorporating adaptive evolution (Rasmussen and Stadler 2019). Our approach also assumes infinite sites and the absence of homoplasies. While these assumptions are limiting over longer periods of sequence evolution, our approach is intended to be used for emerging viral pathogens, sampled over shorter periods of time, when levels of genetic diversity are still low. As such, these assumptions will likely not be violated in cases where this approach will come in useful.

The analysis we presented here focuses on statistical inference using sequence data alone. In recent years, there has also been a growing interest in combining multiple data sources – for example, sequence data and epidemiological data or serological data - to more effectively estimate model parameters. The few studies that have managed to incorporate additional data while performing phylodynamic inference have shown the value in pursuing this goal (Rasmussen et al. 2011; Li et al. 2017). As a next step, we aim to extend the segregating sites approach developed here to incorporate epidemiological data and/or serological data more explicitly. Straightforward extension is possible due to the state-space model structure that is at the core of the particle filtering routine we use.

Our analysis focused on phylodynamic inference based on sequence data belonging to a single viral lineage, with either a single index case or multiple introductions from an outside reservoir. Our approach, however, can be expanded in a straightforward manner to multiple viral lineages. This is especially useful in cases like SARS-CoV-2, where many regions have witnessed the introduction of multiple clades, each fueling the start of more local epidemics (Gonzalez-Reiche et al. 2020; Miller et al. 2020). In this case, a single segregating sites trajectory would be calculated for each clade, such that multiple segregating site trajectories could be fit to at the same time, under specified constraints such as the basic reproduction number being the same across all clades or specified sets of clades. When distinct sets of clades are allowed to differ in their reproductive numbers, questions relating to the selective advantage of some clades over others can also be addressed. As such, this inference method, designed for emerging pathogens with low levels of genetic diversity, may continue to be useful for endemic pathogens when questions involving emerging clades are a focus.

## Materials and Methods

Epidemiological model simulations and calculation of segregating site trajectories. To generate simulated (mock) data, as well as for inference purposes, we specify a compartmental epidemiological model (of arbitrary complexity) and simulate the model under demographic stochasticity using Gillespie’s *τ*-leap algorithm. As a concrete example of a compartmental epidemiological model, we here use a susceptible-exposed-infected-recovered (SEIR) model whose stochastic dynamics are governed by the following equations:

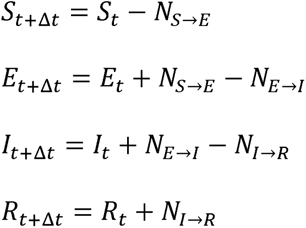

where:

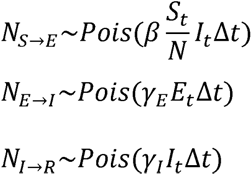

Here, *β* is the transmission rate, N is the host population size, *γ*_E_ is the rate of transitioning from the exposed to the infected class, *γ*_I_ is the rate of recovering from infection, and *Δ*t is the *τ*-leap time step used. R_0_ is given by *β* /*γ*_I_. The epidemiological dynamics of this model can be simulated from the above equations alone. Additional complexity is needed to incorporate virus evolution throughout the time period of the simulation. To incorporate virus evolution, we partition exposed individuals and infected individuals into genotype classes, with basal genotype (genotype 0) being the reference genotype present at the start of the simulation. Mutations to the virus occur at the time of transmission, with the number of mutations that occur in a single transmission event given by a Poisson random variable with mean *μ*, the per-genome, per-transmission event mutation rate. We assume infinite sites such that new mutations necessarily result in new genotypes. New genotypes are numbered chronologically according to their appearance. When new mutations are generated at a transmission event, the new genotype harbors the same mutation(s) as its parent genotype plus any new mutations, which are similarly numbered chronologically based on appearance. We use a sparse matrix approach to store genotypes and their associated mutations to save on memory.

There are three types of events that occur in the SEIR model simulations: transitions from exposed to infected; transitions from infected to recovered; and transmission. To simulate transitions from exposed to infected, during a time step *Δ*t, *N_E_*_→_*_I_* individuals are drawn at random from the set of individuals who are currently exposed. These individuals will transition to the infected class during this time step, while retaining their current genotype statuses. To simulate transitions from infected to recovered, during a time step *Δ*t, *N_E_*_→_*_R_* individuals are drawn at random from the set of individuals who are currently infected. These individuals will transition to the recovered class during this time step. To simulate transmission, during a time step *Δ*t, we add *N_S_*_→_*_E_* new individuals to the set of exposed individuals. For each newly exposed individual, we randomly choose (with replacement) a currently infected individual as its ‘parent’. If no mutations occur during transmission, then this new individual enters the same genotype class as its parent. If one or more mutations occur during transmission, this new individual enters a new genotype class, and the sparse matrix is extended to document the new genotype and its associated mutations (given as integers, without a bitstring or explicit genome structure).

We start the simulation with one infected individual carrying a viral genotype that we consider as the ‘reference’ genotype (genotype 0). To calculate a time series of segregating sites, we define a time window length T (T > *Δ*t) of a certain number of days and partition the simulation time course into discrete, non-overlapping time windows. During simulation, we keep track of the individuals that recover (transition from I to R) within a time window. For each time window i, we then sample n_i_ of these individuals at random, where n_i_ is the number of sequences sampled in a given time window based on the sampling scheme chosen. Because we have the genotypes of the sampled individuals from the sparse matrix, we can calculate for any time window i, the number of segregating sites S_i_. S_i_ is simply the number of polymorphic sites across the sampled individuals in time window i. While in our simulations, we sample individuals as they recover, alternative sampling schemes can instead be assumed. For example, individuals could be sampled as they transition from the exposed to the infected class, or while they are in the infected class.

Epidemiological inference using time series of segregating sites. Our inference approach relies on particle filtering, also known as Sequential Monte Carlo (SMC), to estimate model parameters and reconstruct latent state variables. The underlying forward model we use is formulated as a state-space model, with epidemiological variables (e.g., S, E, I, and R) being latent (that is, unobserved) variables in the process model. The model is simulated using Gillespie’s *τ*-leap algorithm, as described in the section above. The evolutionary component of the model also contributes to the process model. For the observation model, we perform k ‘grabs’ of sampled individuals, with each grab consisting of the following steps:

- pick (without replacement) n_i_ individuals from the set of individuals who recovered during time window i, where n_i_ is the number of samples present in the empirical dataset in window i. We sample the same number of individuals as in the segregating sites dataset that the model interfaces with, since sampling effort impacts the number of segregating sites.
- calculate the simulated number of segregating sites S ^sim^, based on the genotypes of the sampled n_i_ individuals.

Between grabs, replacement of previously sampled individuals occurs. We then calculate the mean number of segregating sites for window i by taking the average of all k S ^sim^ values. Finally, we calculate the probability of observing S_i_ segregating sites in window i, given the model-simulated mean number of segregating sites, using a Poisson probability mass function parameterized with the mean S ^sim^ value and evaluated at S_i_. These probabilities serve as the weights for the particles. Particle weights are calculated at the end of each time window with n_i_ > 0. Particles are resampled at the end of each of these time windows according to their assigned weights. Particles with stochastic extinction of the virus prior to the end of the last time window with n_i_ > 0 have weights set to 0 in time window i. If the number of sampled individuals n_i_ in time window i exceeds the total number of individuals who recovered in time window i, the particle weight is similarly set to 0.

Latent state variables are reconstructed by randomly sampling a particle at the end of an SMC simulation and plotting the values of its simulated latent state variables over time. All of our SMC simulations were performed with 200 particles and k = 50 grabs. For inference on the mock dataset, a population size of 100,000 was used, and we set the mean duration of the exposed period to be 2 days (1/*γ*_E_) and the mean duration of the infectious period to be 3 days (1/*γ*_I_).

Note that the complexity of this inference method is largely independent of the number of input sequences. This stands in contrast to phylodynamic inference approaches that frequently downsample sequences to reduce runtime.

Implementation of the transmission heterogeneity model. We implement transmission heterogeneity by splitting the infected classes into a high-transmission and a low-transmission class, as has been done elsewhere (Volz and Siveroni 2018; Miller et al. 2020). For an SEIR model, the model extended to incorporate transmission heterogeneity becomes:

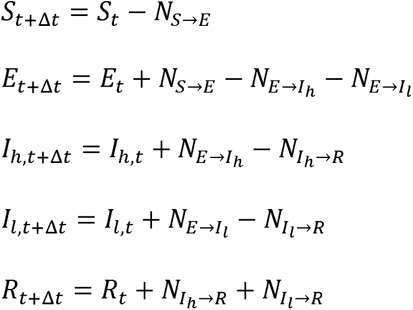

where:

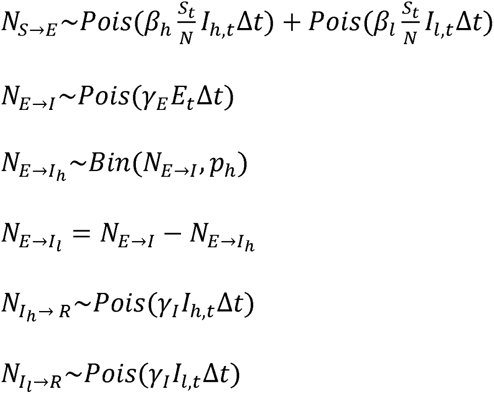

The parameter p_h_ quantifies the proportion of exposed individuals who transition to the high- transmission I_h_ class. Parameters *β*_h_ and *β*_l_ quantify the transmission rates of the infectious classes that have high and low transmissibility, respectively. We set the values of *β*_h_ and *β*_l_ based on a given parameterization of overall R_0_ and the parameter p_h_. To do this, we first define, as in previous work (Volz and Siveroni 2018; Miller et al. 2020), the relative transmissibility of infected individuals in the I_h_ and I_l_ classes as 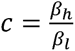. We further define a parameter P as the fraction of secondary infections that result from a fraction p_h_ of the most transmissible infected individuals. Based on given values of p_h_ and P, we set c, as in previous work (Miller et al. 2020), to 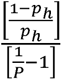. With c defined in this way, p_h_ is interpreted as the proportion of most infectious individuals that result in P of secondary infections. We set P to 0.80, to make p_h_ easily interpretable relative to the “20/80” rule in disease ecology (Woolhouse et al. 1997). Recognizing that 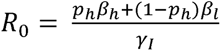 in this model, we can then solve for *β*: 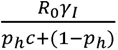 and set *β_h_ = cβ_l_*.

Converting simulated sequences into nucleotide sequences for the performance comparison against PhyDyn. Simulated sparse matrices were converted to nucleotide alignments by first generating a reference sequence with the same length as the maximum number of mutations in the sparse matrix and choosing an A,C,G, or T nucleotide at each site with equal probability. A mutated sequence was generated for each genotype represented in the sparse matrix by replacing the reference allele at that position with another nucleotide chosen with equal probability. The final FASTA alignment was generated by identifying the simulated sequence associated with each sampled individual. Generation of the simulated FASTA file was done using Python v3.9.4 with Numpy v1.19.4.

The simulated FASTA alignment was used to generate a BEAST2 XML file from template XML which was generated in part using Beauti v2.6.6. This template used a JC69 nucleotide substitution model with no invariant sites. We assumed an uncorrelated log-normally distributed relaxed clock with a uniform [0.0, 1E-2] prior on the mean and a uniform [0.0,2.0] prior on the standard deviation.

A single-deme structured coalescent prior as defined by the following equations was implemented using PhyDyn v1.3.8. All sampled sequences were assigned to the “I” class:

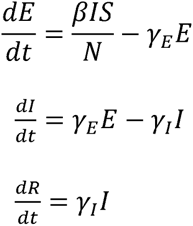

where: (3 == R_o_*γ_I_*. A population size of 100,000 with a single initially infected individual was used. We assume infected individuals remain exposed for an average of 2 days (1/*γ*_E_) and infectious (1/*γ*_I_) for an average of 3 days. R_0_ was estimated using a uniform [1.0, 10.0] prior.

Sampled parameters and trees were logged every 1000 states and all MCMC chains were run for at least 90M iterations. The first 10% of MCMC chains were discarded as burn-in and the ESS values of all parameters were >200, as identified by Tracer v1.7.1 (10.1093/sysbio/syy032).

Epidemiological model structure and parameterization used for the SARS-CoV-2 analysis. The process model we use in our application to SARS-CoV-2 sequence data from France is based on a previously published epidemiological model (Salje et al. 2020). We base our process model on this published model to allow for a direct comparison of inferred R_0_ values between our sequence-based analysis and their analysis that focuses on SARS-CoV-2 spread in France over a similar time period. Their analysis was based on fitting an epidemiological model to a combination of case, hospitalization, and death data. Their model structure, once implemented using Gillespie’s *τ*-leap algorithm, is given by:

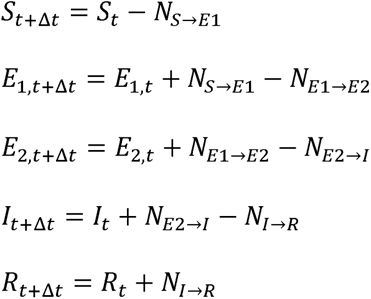

where

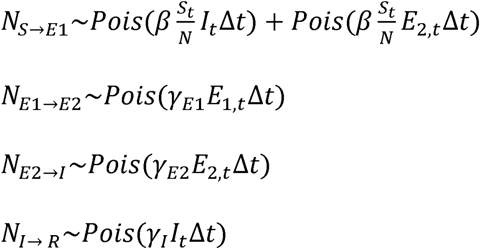

with *β* being the transmission rate, the average duration of time spent in the E_1_ class given by 1/y_E1_ = 4 days, the average duration of time spent in the E_2_ class given by 1/y_E2_ = 1 day, and the average duration of time spent in the infected class given by 1/*γ_I_* = 3 days. Their model assumes that the transmission efficiency *β* of exposed class 2 (E_2_) and that of the infected class *I* are the same; their model considers E_2_ and I as distinct classes to interface with the case data, where symptoms are assumed to not appear before an individual is infected (in class I). We maintain the model structure with E_1_, E_2_, and I rather than reducing it to a model structure with just a single E and a single I class to keep the same distribution of infection times as in their model.

Because SARS-CoV-2 dynamics are characterized by substantial levels of transmission heterogeneity (Adam et al. 2020; Miller et al. 2020; Sun et al. 2021) and we have shown in Figure 1 that transmission heterogeneity impacts segregating site trajectories, we expanded the compartmental epidemiological model described above to include transmission heterogeneity in a manner similar to the one we used in Figure 1. Based specifically on the analysis by Paireau and colleagues (Paireau et al. 2022), we set p_h_ to 0.10, such that 10% of infections are responsible for 80% of secondary infections. Analogous to the approach we undertook for the simulated data, we estimated R_0_ and t_0_ using the segregating site trajectory shown in Figure 6B.

Based on phylogenetic analyses that have indicated that early introductions of SARS-CoV-2 into focal regions likely resulted from multiple introductions rather than a single one, we further considered a modified version of the epidemiological model that would allow multiple introductions. The modification relied on the incorporation of infections that resulted from contact with an outside-of-France viral reservoir. Similar to the approach adopted by some existing phylodynamic analyses, e.g., (Geidelberg et al. 2021), the viral population dynamics in this reservoir are simplified to exponential growth. This infected population from outside of France acts as another source of infection for susceptible individuals within France, allowing the multiple introductions of SARS-CoV-2 into France.

As in the focal region, new genotypes are expected to emerge in the outside reservoir. As we assume an infinite sites model, the genotypes that emerge in the outside reservoir and in the focal region will not overlap except in the basal genotype that is first introduced to the focal region. For this reason, and because the basal genotype is expected to be more prevalent than any of the viral genotypes that stem from it, we consider only the repeated introduction of the basal genotype into France. Starting at the time of emergence of the basal genotype in the outside reservoir, we let the number of individuals infected with this basal genotype Y_t_ grow exponentially:

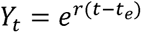

where r is the intrinsic growth rate of the basal genotype and t_e_ is the time of emergence for the basal genotype. We set the intrinsic growth rate to 0.22 day^-1^, based on empirical estimates (Dehning et al. 2020; Musa et al. 2020). To set t_e_, we first identified the genotype sampled in France that is genetically closest to the reference strain Wuhan/Hu-1 (MN908947.3). This basal genotype differs from Wuhan/Hu-1 by 4 nucleotides: C241T, C3037T, C14408T, and A23403G. Using GISAID data, we then identified sequences with collection locations outside of France that carried all four of these mutations that define the basal genotype. The earliest of these sequences including the four basal genotype-defining mutations was collected on January 8^th^, 2020 in the Netherlands, suggesting that the basal genotype had been circulating prior to January 8^th,^ 2020. Considering the potential delay between emergence and the time of first detection, we considered three distinct t_"_ values: December 24^th^, 2019, January 1^st^, 2020, and January 8^th^, 2020.

Individuals infected in this outside reservoir can transmit their infection to susceptible individuals within France. The rate at which they transmit the infection is reduced relative to the rate at which infected individuals within France transmit infection to susceptible individuals within France. We let the factor by which transmission is reduced be given by the factor *η*.

During a *τ*-leap timestep, the number of individuals within France who become infected from contact with an infected individual outside of France is therefore given by:

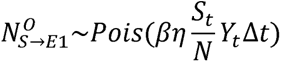

As we are considering only the dynamics of the basal genotype in the outside reservoir, all of these newly infected individuals will carry the basal genotype unless mutation occurs during the transmission process.

### Estimation of the per genome, per transmission event mutation rate

We set the per-genome, per-transmission mutation rate parameter *μ* to 0.33. This is based on the fit of a Poisson distribution to the number of de novo substitutions that were observed in 87 transmission pairs of SARS-CoV-2 from four studies (James et al. 2020; Popa et al. 2020; Braun et al. 2021; Lythgoe et al. 2021). Accession numbers for 78/87 of these transmission pairs are available in Table S1. Accession numbers for the remaining pairs were provided by the corresponding authors of the relevant publication (Lythgoe et al. 2021). Sequence data were aligned to Wuhan/Hu-1 (MN908947.3) (Wu et al. 2020) using MAFFT v.7.464 (Katoh 2002). Insertions relative to Wuhan/Hu-1 were removed and the first 55 and last 100 nucleotides of the genome were masked. De novo substitutions for each pair were identified in Python v.3.9.4 (http://www.python.org) using NumPy v.1.19.4 (Harris et al. 2020). Ambiguous nucleotides were considered in the identification of de novo substitutions (i.e. an R nucleotide was assumed to match both an A and a G). The mean number of substitutions between transmission pairs is the Maximum Likelihood Estimate for the λ parameter of the Poisson distribution. The 95% confidence intervals were calculated using the exact method using SciPy v.1.5.4 (SciPy 1.0 Contributors et al. 2020) such that the lower bound was 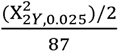 and the upper bound was 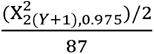 where Y is the total number of observed substitutions.

The value for *μ* = 0.33 is consistent with population-level substitution rate estimates for SARS-CoV-2, which range from 7.9 x 10^-4^ to 1.1 x 10^-3^ substitutions per site per year (Duchene et al. 2020; Pekar et al. 2020). With a genome length of SARS-CoV-2 of approximately 30,000 nucleotides and a generation interval of approximately 4.5 days (Griffin et al. 2020), these population-level substitution rates would correspond to per genome, per transmission mutation rates of between 0.29 and 0.41, respectively.

Estimation of segregating site trajectories for the France data.

We downloaded all complete and high-coverage SARS-CoV-2 sequences with complete sampling dates sampled through March 17^th^, 2020 (https://www.france24.com/en/20200316-live-france-s-macron-addresses-nation-amid-worsening-coronavirus-outbreak) in France and uploaded through April 29^th^, 2021 from GISAID (Shu and McCauley 2017). Sequences were aligned to Wuhan/Hu-1 using MAFFT v.7.464 Insertions relative to Wuhan/Hu-1 were removed. Any sequences with fewer than 28000 A, C, T, or G characters were removed. Following this filtering protocol our dataset included 479 sequences. We masked the first 55 and last 100 nucleotides in the genome as well as positions marked as “highly homoplasic” in early SARS-CoV-2 sequencing data (https://github.com/W-L/ProblematicSites_SARS-CoV2/blob/master/archived_vcf/problematic_sites_sarsCov2.2020-05-27.vcf). Pairwise SNP distances were calculated in a manner that accounted for IUPAC ambiguous nucleotides in Python using NumPy. To subset these data to a single clade circulating within France, we identified the connected components of this pairwise distance matrix with a cutoff of 1 SNP in Python using SciPy and identified the shared SNPs relative to Wuhan/Hu-1 between all sequences in each connected component. The largest connected component contained 308 sequences which shared the substitutions C241T, C3037T, C14408T, and A23403G. Our final dataset included these 308 as well as 122 sequences from connected components that shared these four substitutions relative to Wuhan/Hu-1. We included connected components in which all sequences had an N at any of the four clade-defining sites of the largest connected component. Two sequences were excluded as they differed from all other sequences in the dataset by > 7 SNPs. This dataset is similar to the set of sequences analyzed in Danesh et al. (2020). Sequences were binned into four-day windows, aligned such that the last window ended on the latest sampling date. The number of segregating sites in each window was calculated in Python using NumPy. Ambiguous nucleotides were considered in the calculation of segregating sites.

### Phylogenetic analysis of SARS-CoV-2 sequences from France

To confirm that the subset of sequences from France obtained from finding connected components formed an evolutionary lineage/clade, we first combined the 479 sequences sampled from France with 100 randomly-selected complete, high-coverage, collected date complete sequences sampled from outside France through March 17^th^, 2020 and uploaded to GISAID through April 29^th^, 2021. These sequences were aligned to Wuhan/Hu-1 using MAFFT, insertions were removed, and the sites described above were masked. This alignment was concatenated with the aligned sequences from France. IQ-Tree v. 2.0.7 (Minh et al. 2020) was used to construct a maximum likelihood phylogeny, and ModelFinder (Kalyaanamoorthy et al. 2017) was used to find the best fit nucleotide substitution model (GTR+F+I). Small branches were collapsed. TreeTime v. 0.8.0 (Sagulenko et al. 2017) was used to remove any sequences with more than four interquartile distances from the expected evolutionary rate, rooting at Wuhan/Hu-1. Treetime was also used to generate a time-aligned phylogeny assuming a clock rate of 1 x 10^-3^ with a standard deviation of 5 x 10^-4^, a skyline coalescent model, marginal time reconstruction, accounting for covariation, and resolving polytomies.

Maximum likelihood phylogenies were visualized in Python using Matplotlib v. 3.3.3 (Hunter 2007) and Baltic (https://github.com/evogytis/baltic).

### Availability of code

Python code used for generation of all figures is available on GitHub: https://github.com/koellelab/segregating-sites

---

**Figure S1.**
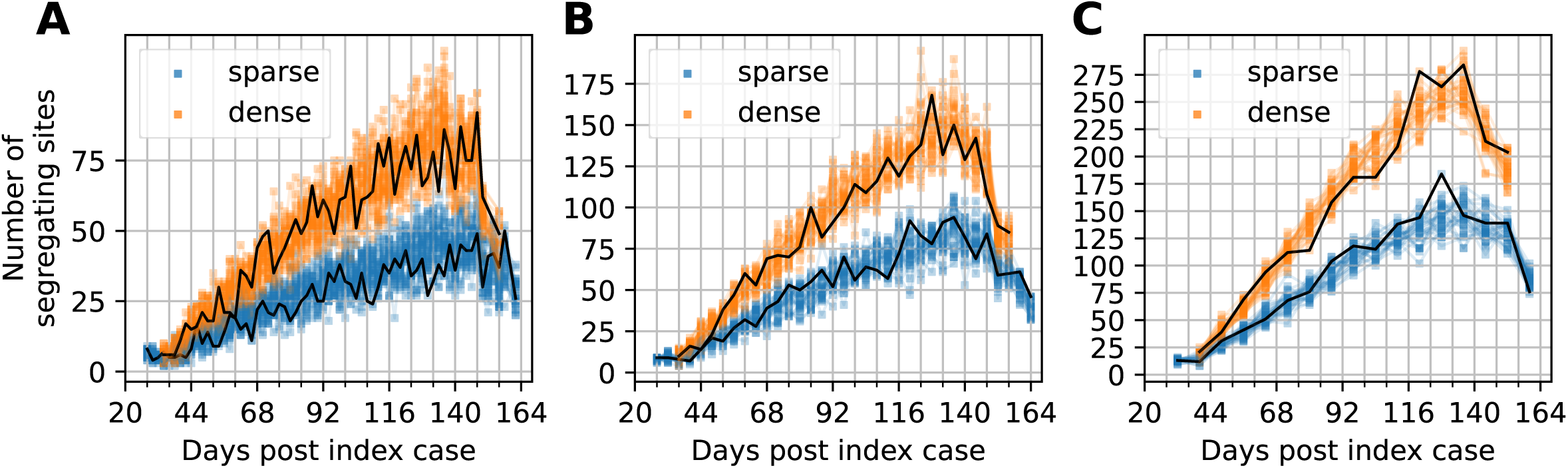
Segregating site trajectories under different time window lengths. Segregating site trajectories for the simulation shown in Figure 1A under dense (orange) and sparse (blue) sampling effort, when trajectories are calculated using time window lengths of (A) 2 days; (B) 4 days; and (C) 6 days. Under the dense sampling scheme, sampling efforts are 20 sequences per 2 day time window (A), 40 sequences per 4 day time window (B), and 60 sequences per 6 day time window (C). Under the sparse sampling scheme, sampling efforts are 10 sequences per 2 day time window (A), 20 sequences per 4 day time window (B), and 30 sequences per 6 day time window (C). 30 randomly-sampled segregating site trajectories are shown for each sampling effort. Black lines show a single representative segregating site trajectory under the dense and sparse sampling efforts, respectively. Shorter time window lengths yield segregating site trajectories that are more variable between time windows, while longer time window lengths decrease the resolution on temporal trends in the number of segregating sites. Shorter time window lengths also result in a lower number of segregating sites, reflecting the smaller number of sequences per time window.

**Figure S2.**
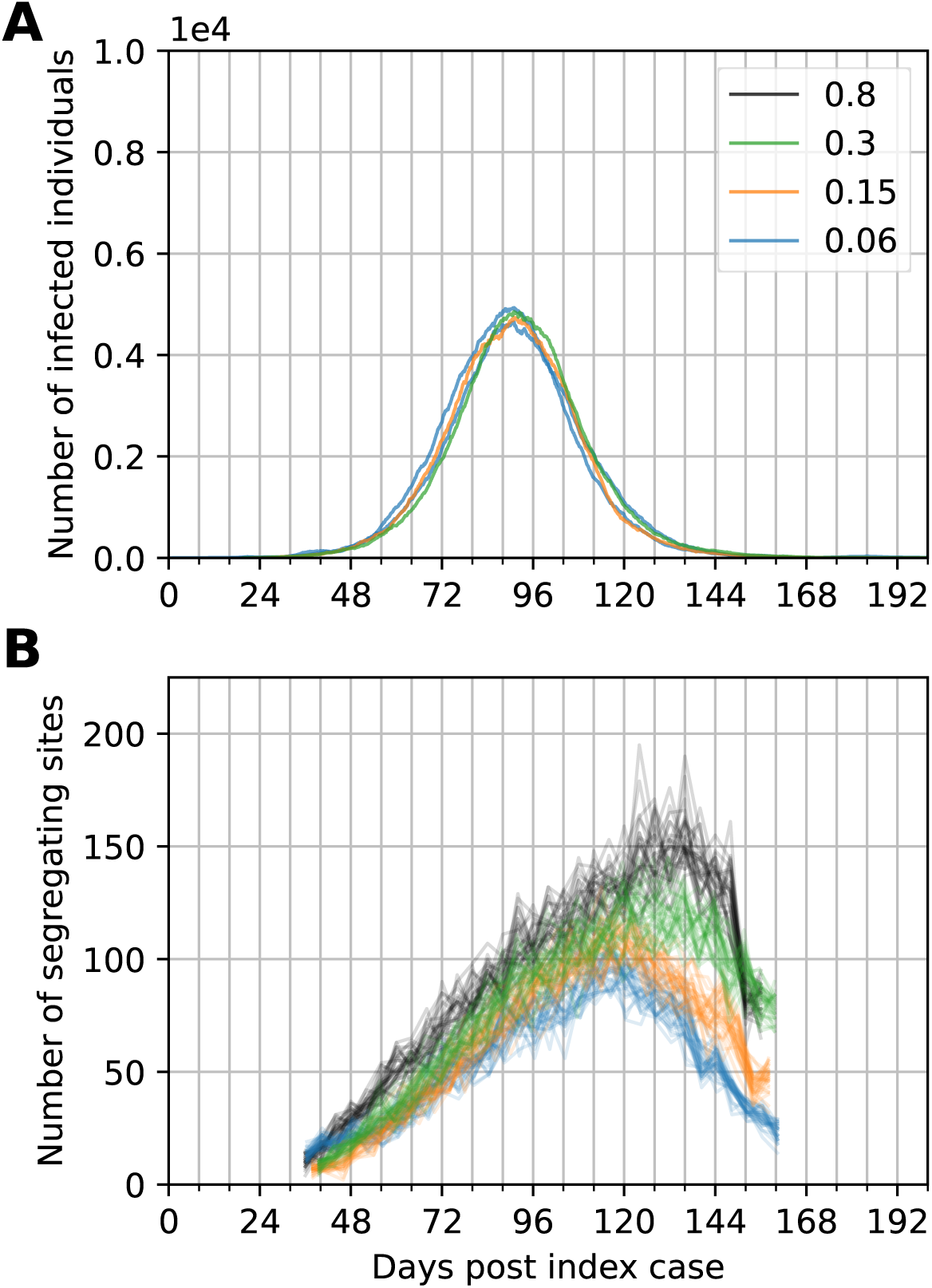
Segregating site trajectories under different levels of transmission heterogeneity. (A) Simulated dynamics of infected individuals (I) under an SEIR model with an R_0_ of 1.6 and incorporating various levels of transmission heterogeneity compared to those of the original R_0_ = 1.6 simulation without transmission heterogeneity. Transmission heterogeneity simulations shown are all shifted in time to align their epidemic peaks with the simulation without transmission heterogeneity (blue line; p_h_ = 0.8). The transmission heterogeneity simulations considered span from low levels of transmission heterogeneity (p_h_ = 0.3, corresponding to 30% of the most infectious individuals responsible for 80% of secondary infections), to higher levels (p_h_ = 0.06, corresponding to 6% of the most infectious individuals responsible for 80% of secondary infections. Intermediate levels of transmission heterogeneity are parameterized with p_h_ = 0.15, corresponding to 15% of the most infectious individuals responsible for 80% of secondary infections. (B) Segregating site trajectories for the simulations shown in (A). All simulations are densely sampled (40 sequences sampled per time window of 4 days).

**Figure S3.**
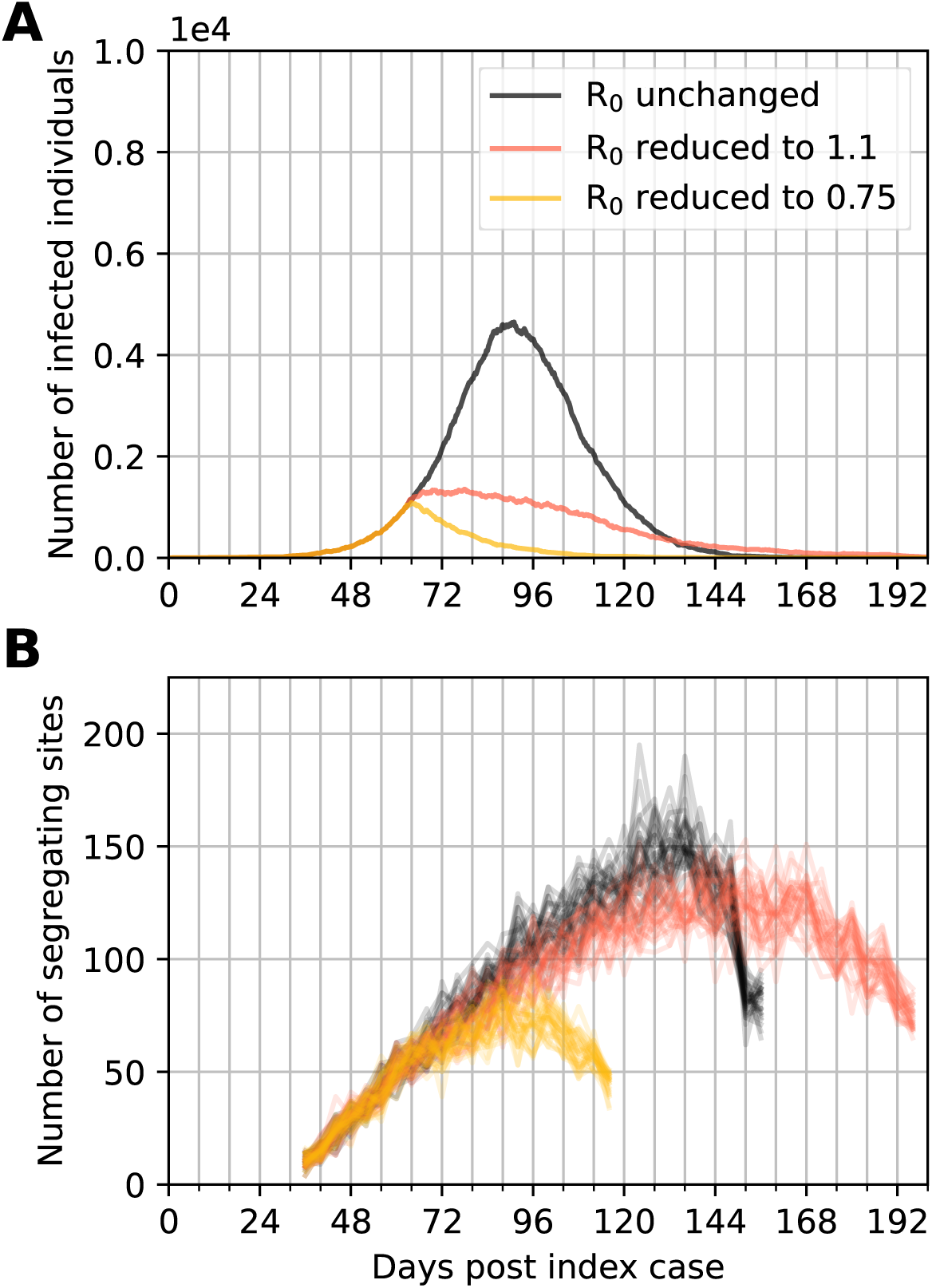
Segregating site trajectories under transmission reduction scenarios occurring at a later time point of the epidemic. (A) Simulated dynamics of infected individuals (I) under an SEIR model simulated with changing R_0_. Changes in R_0_ occurred when the number of infected individuals reached 1000. The simulation in red has R_0_ decreasing to 1.1. The simulation in yellow has R_0_ decreasing to 0.75. The simulation in blue has R_0_ remaining at 1.6. (B) Segregating site trajectories for the three simulations shown in (A).

**Figure S4.**
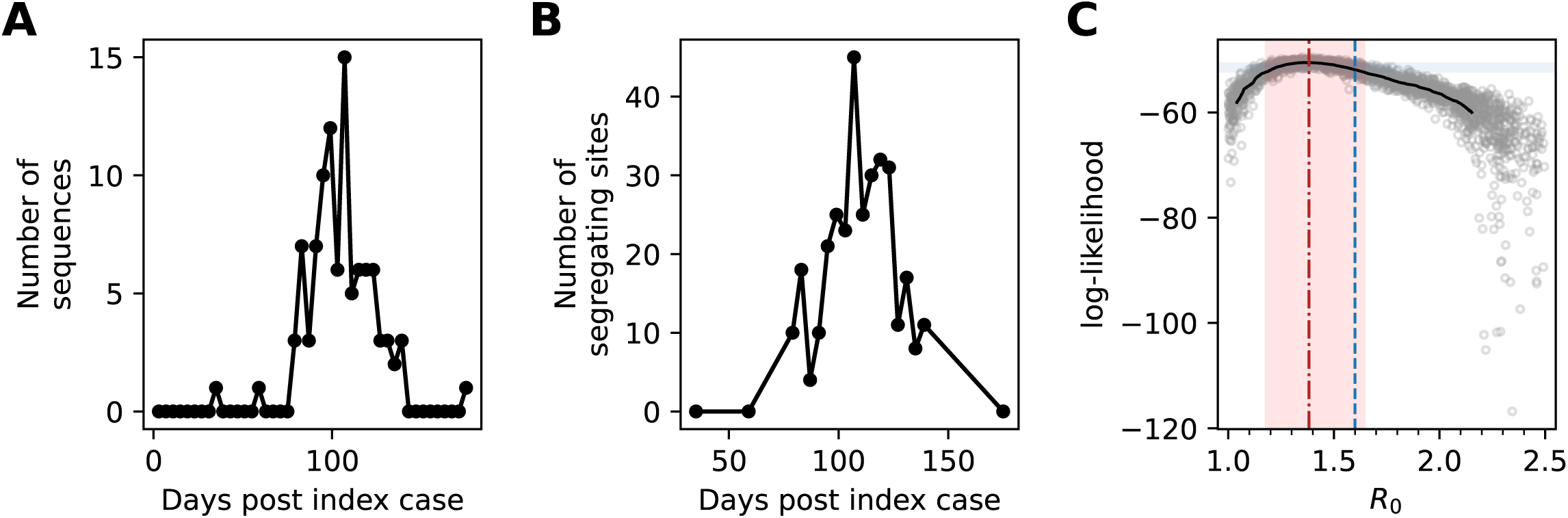
Epidemiological inference on a simulated trajectory of segregating sites, with lower sampling effort than in Figure 2. (A) The number of sampled sequences over time, by time window. Sampling was done in proportion to the number of individuals recovering in a time window. In all, 100 sequences were sampled over the course of the simulated epidemic. (B) Simulated segregating site trajectory from the sampled sequences. (C) Estimation of R_0_ using SMC. The maximum likelihood estimate for R_0_ was 1.38 [95% CI = 1.17 to 1.65].

**Figure S5.**
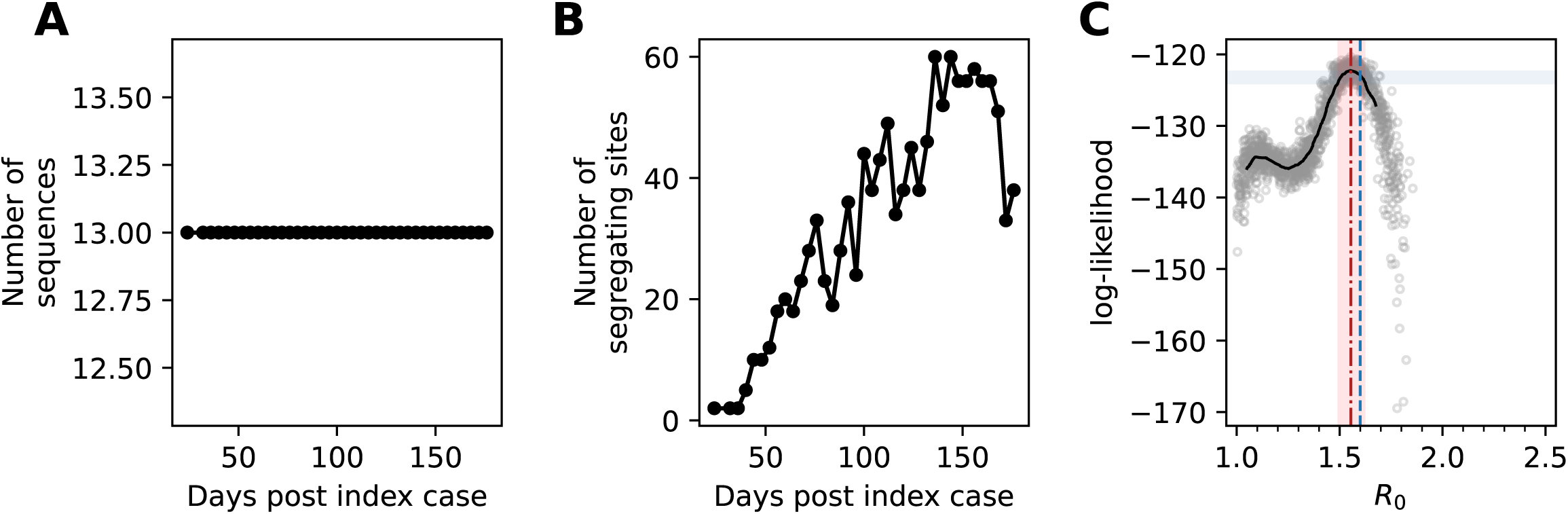
Epidemiological inference on a simulated trajectory of segregating sites, with uniform rather than proportional sampling. (A) The number of sampled sequences, by 4-day time window. Uniform sampling was performed by sampling 13 sequences per time window. Time windows with fewer than 13 sequences available were not included in the analysis. As such, here, only 494 sequences were used for inference. (B) Simulated segregating site trajectory from the sampled sequences. (C) Estimation of R_0_ using SMC. The estimate for R_0_ was 1.55 [95% CI = 1.49 to 1.63]. In contrast, the proportional sampling strategy shown in Figure 2 had a higher level of uncertainty in its R_0_ estimate (1. 54, with 95% CI = 1.37 to 1.81).

**Figure S6.**
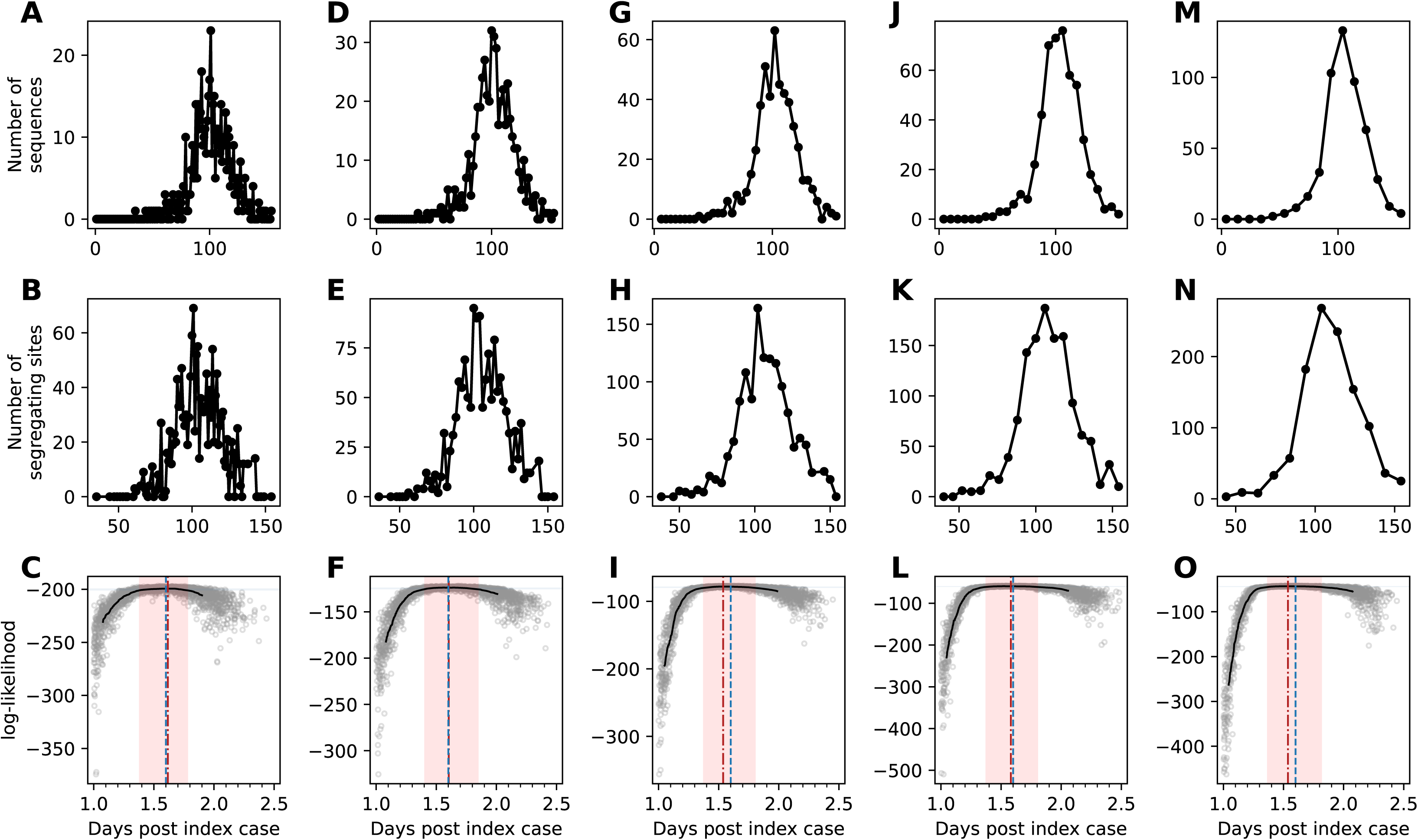
Epidemiological inference on the same set of sampled sequences as in Figure 2, binned at different time window lengths. For all time window lengths, the same set of 500 proportionally- sampled sequences were used. Top row: The number of sampled sequences over time, binned by time window (1, 2, 4, 6, and 10 days, respectively). Middle row: Segregating site trajectories from the set of binned sequences. Bottom row: Estimation of R_0_ using SMC. Panels A-C show results using a time window length of 1 day. The estimate for R_0_ was 1.62 [95% CI = 1.38 to 1.78]. Panels D-F show results using a time window length of 2 days. The estimate for R_0_ was 1.60 [95% CI = 1.40 to 1.85]. Panels G-I reproduce the results shown in Figure 2, using a time window length of 4 days. The estimate for R_0_ was 1.54 [95% CI =1.37 to 1.81]. Panels J-L show results using a time window length of 6 days. The estimate for R_0_ was 1.58 [95% CI = 1.37 to 1.81]. Panels M-O show results using a time window length of 10 days. The estimate for R_0_ was 1.54 [95% CI = 1.36 to 1.82].

**Figure S7.**
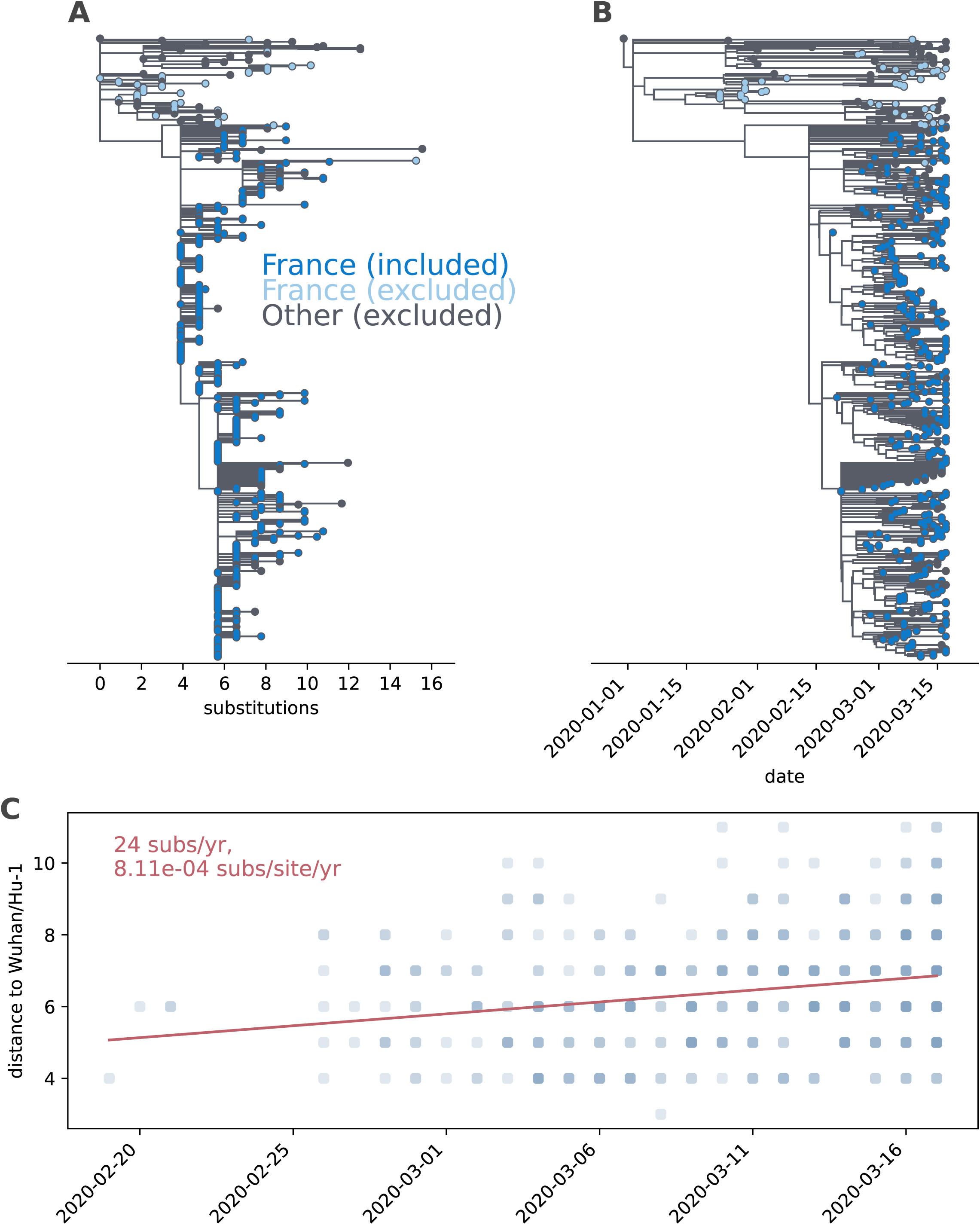
Inferred phylogenies for the sequences sampled from France, January 23-March 17, 2020. (A) Divergence tree, showing the number of nucleotide substitutions from Wuhan/Hu-1. Sequences from France are colored in blue, with dark blue coloring indicating sequences that were included in our single-lineage analysis and light blue coloring indicating sequences that were excluded from our analysis. Tips colored in gray denote genetically similar sequences sampled from outside of France during this time period. (B) Time-aligned maximum likelihood phylogeny, with coloring of sequences as in (A).

Table S1. Transmission pairs used to estimate the per-genome, per-transmission event mutation rate *μ*. Accession numbers of the consensus sequences from the donor and the recipient of the transmission pair are provided.

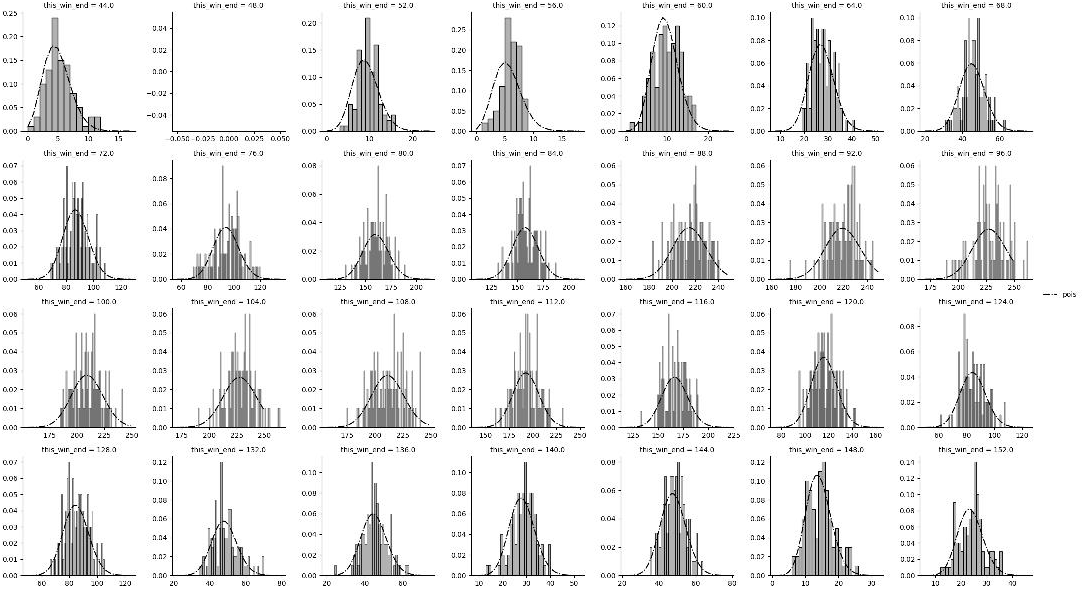

